# Sensory plasticity of dorsal horn silent neurons: a critical mechanism for neuropathic pain

**DOI:** 10.1101/2025.09.26.678726

**Authors:** Ahmed Negm, François Gabrielli, Mickael Zbili, Louis Poncet-Megemont, Karine Herault, Jennifer Makhoul, Radhouane Dallel, Cedric Peirs

## Abstract

The spinal cord dorsal horn (DH) integrates and modulates sensory processing but undergoes critical plasticity following nerve injury, leading to pain hypersensitivity. Mechanical allodynia, or touch-evoked pain, is a highly prevalent and debilitating symptom of neuropathic pain. It has been proposed that, after nerve injury, innocuous sensory neurons gain access to nociceptive-specific (NS) circuits in the DH due to altered spinal inhibitory controls, thereby converting touch into pain. It is however unclear how sensory processing is reorganized in these conditions across the different laminae of the DH to generate this symptom.

In this study, we developed a novel *ex vivo* somatosensory preparation to selectively analyze excitatory neuronal activity across all DH laminae simultaneously, following physiological stimulations of the skin. Using two-photon calcium (Ca^2+^) imaging, we studied the DH activity under physiological conditions, after spinal disinhibition or nerve injury, and generated a computational model to reveal the sensory plasticity of individual DH neurons that leads to neuropathic pain.

We demonstrate that spinal disinhibition, whether pharmacologically induced or resulting from nerve injury, converts most DH excitatory neurons into highly polymodal cells. We further show that such disinhibition unmasks an unprecedented number of previously silent neurons in both superficial and deep DH laminae, responding to a wide dynamic range (WDR) of sensory modalities. The computational model pinpoints that neuropathic pain does not result primarily from the transformation of excitatory NS neurons into WDR neurons, but rather from the activation of a previously dormant excitatory circuit. This newly active circuit spans both superficial and deep DH laminae and is predominantly composed of WDR excitatory neurons The identification of this extensive silent neuronal network provides critical insights into DH plasticity mechanisms underlying neuropathic pain, and should guide future therapeutic strategies.

**Highlights:** Sensory modalities of dorsal horn neurons are defined by spinal inhibition

Neuropathic mechanical allodynia does not result from the transformation of nociceptive specific neurons into wide dynamic range neurons

Neuropathic pain is mediated by the activation of a previously silent circuit

## INTRODUCTION

Unlike acute pain, which serves to protect against injury, neuropathic pain is a debilitating condition affecting 7-10% of the population worldwide (Bouhassira, 2019). Defined as pain caused by a lesion or disease of the somatosensory nervous system, neuropathic pain is characterized by its long-term nature and resistance to most therapeutic interventions (Attal and Bouhassira, 2021). The dorsal horn (DH) of the spinal cord is the first central relay that processes sensory information from peripheral tissues, and a major site of central sensitization that leads to neuropathic pain (von Hehn et al., 2012). Nerve injury induces a cascade of molecular and cellular changes in the DH, resulting in sensitization of DH neurons and pain hypersensitivity. While various mechanisms and neuronal populations have been proposed to contribute to central sensitization (Peirs and Seal, 2016), there is consensus that these processes disrupt the balance between excitation and inhibition in the DH, resulting in an aberrant activation of DH sensory neurons. Spinal inhibition plays a crucial role in the plasticity of sensory processing under pathological conditions, as patients with genetic mutations affecting their glycinergic signaling suffer from exaggerated pain (Vuilleumier et al., 2018), and spinal disinhibition invariably leads to symptoms of neuropathy in preclinical models (Gradwell et al., 2019). To date, most studies converge on the idea that innocuous and noxious neuronal circuits are normally segregated at the level of the DH, through the protective control of inhibitory interneurons. After nerve injury, these inhibitory controls would be reduced, leading to the activation of nociceptive specific neurons (NS) in superficial laminae by low-threshold mechanosensitive (LTM) neurons from the deep DH, thus turning touch into pain. However, activation of NS neurons by LTM afferents as a main mechanism of neuropathic pain remains controversial (Andrew, 2009; Chapman et al., 1998; Keller et al., 2007; Lavertu et al., 2014), and selective inhibition or ablation of lamina I DH neurons, including those that belong to the major NS spino-thalamic (STT) ascending pathway, has failed to abolish pain hypersensitivity in pathological conditions (Huang et al., 2019; Maiaru et al., 2018; Mantyh et al., 1997). More recently, a study used the stimulus-coupled transcription technology CANE (capturating activated neuronal ensembles) and showed that silencing NS neurons did not relieve the hypersensitivity from neuropathic mice to any sensory stimulus, challenging the role of NS neurons in neuropathic pain and obscuring how activation of LTM sensory neurons leads to pain after nerve injury (Zhang et al., 2024a).

Answering this critical question has been hampered by several factors. First, although *in vivo* electrophysiological methods are best suited to study the function of DH neurons in living tissue, these approaches are often limited to blind recordings, or supervised neuronal sampling. For instance, most of these studies have used mechanical probing to search for receptive field neurons, introducing a sampling bias towards mechano-sensitive neurons. These techniques also favor large over small diameter neurons, which are easier to record from, and use repeated stimulations to search for neurons, which could lead to sensitization of sensory neurons over time and changes of sensory specificity (Wang et al., 2018). In contrast, functional studies using activity markers on postmortem tissues have a better spatial resolution and can more precisely address the phenotype of activated neurons, but are limited in the number of variables that can be tested in each sample (Peirs et al., 2020). Finally, recent works using genetic and laser scanning microscopy have greatly improved temporal, spatial and phenotypic resolution, but overlooked the deep DH that is still out of range of microscopy in living tissue (Barkai et al., 2023; Warwick et al., 2022). A compelling description of which cells are engaged by innocuous stimuli in neuropathic condition is thus still not yet available, especially in the deep DH.

To fill these gaps, we developed a new *ex vivo* somatosensory preparation inspired by two seminal works (Hachisuka et al., 2016; Koerber and Woodbury, 2002). Using 2-photon calcium (Ca^2+^) imaging of a parasagittal DH slice still connected to the skin, we analyzed the sensory modalities of thousands of excitatory neurons across the whole DH, following the stimulation of intact peripheral endings. Unexpectedly, we first found that the majority of DH neurons are completely silenced by spinal inhibition under physiological conditions, in both superficial and deep laminae of the DH. We then showed that most DH neurons receive multiple sensory inputs that are normally finely controlled by spinal inhibition, but which become active after nerve injury. Computational analysis of the sensory plasticity of each individual neurons due to the loss of DH inhibition revealed that the major plasticity of the DH induced by the neuropathy does not lead to the activation of NS neurons by LTM neurons, but to the recruitment of a substantial pool of polymodal and previously silent neurons, which are likely the main drivers of neuropathic pain. This finding challenges the prevailing view that neuropathic mechanical allodynia results from maladaptive activation of nociceptive neurons by innocuous stimuli and emphasizes the importance of defining the molecular identity of these silent neurons. Targeting this largely unknown neuronal population may therefore provide a promising avenue for developing therapies for neuropathic pain, addressing the urgent need to translate mechanistic insights into effective treatments (Finnerup et al., 2021).

## RESULTS

### Characterization of an *ex vivo* parasagittal spinal cord DH – skin somatosensory preparation

Current imaging techniques used to investigate the dynamic of spinal cord circuits are often limited to the most superficial laminae due to the absorption and scattering of light in biological tissues, especially through the abundant white matter that surrounds the DH (Johannssen and Helmchen, 2010). To be able to measure the activity of DH neurons evoked by peripheral physiological stimulations at the circuit level across multiple laminae, we adapted the somatosensory *ex vivo* preparation from (Hachisuka et al., 2016; Koerber and Woodbury, 2002). This new preparation consisted of a parasagittal spinal cord slice still connected to the lumbar dorsal root ganglions (DRG), the tibial branch of the sciatic nerve, and the plantar surface of the hind paw (Fig. 1A). Combining this preparation with 2-photon microscopy, we measured Ca^2+^ transients evoked in the mouse DH from lamina I to V, following several innocuous and noxious stimuli of the skin. Imaging was restricted to excitatory neurons using VGLUT2^Cre/+;^ R26^lsl-GCamp6f/+^ mice, in which conditional expression of the Ca^2+^ sensor GCaMP6f is under the control of the vesicular glutamate transporter 2 promotor (VGLUT2), which is present in virtually all excitatory neurons of the DH (Peirs et al., 2020) (Fig. 1B). A total of 6142 neurons from 37 mice were included in the analysis of this study.

**Figure 1.**
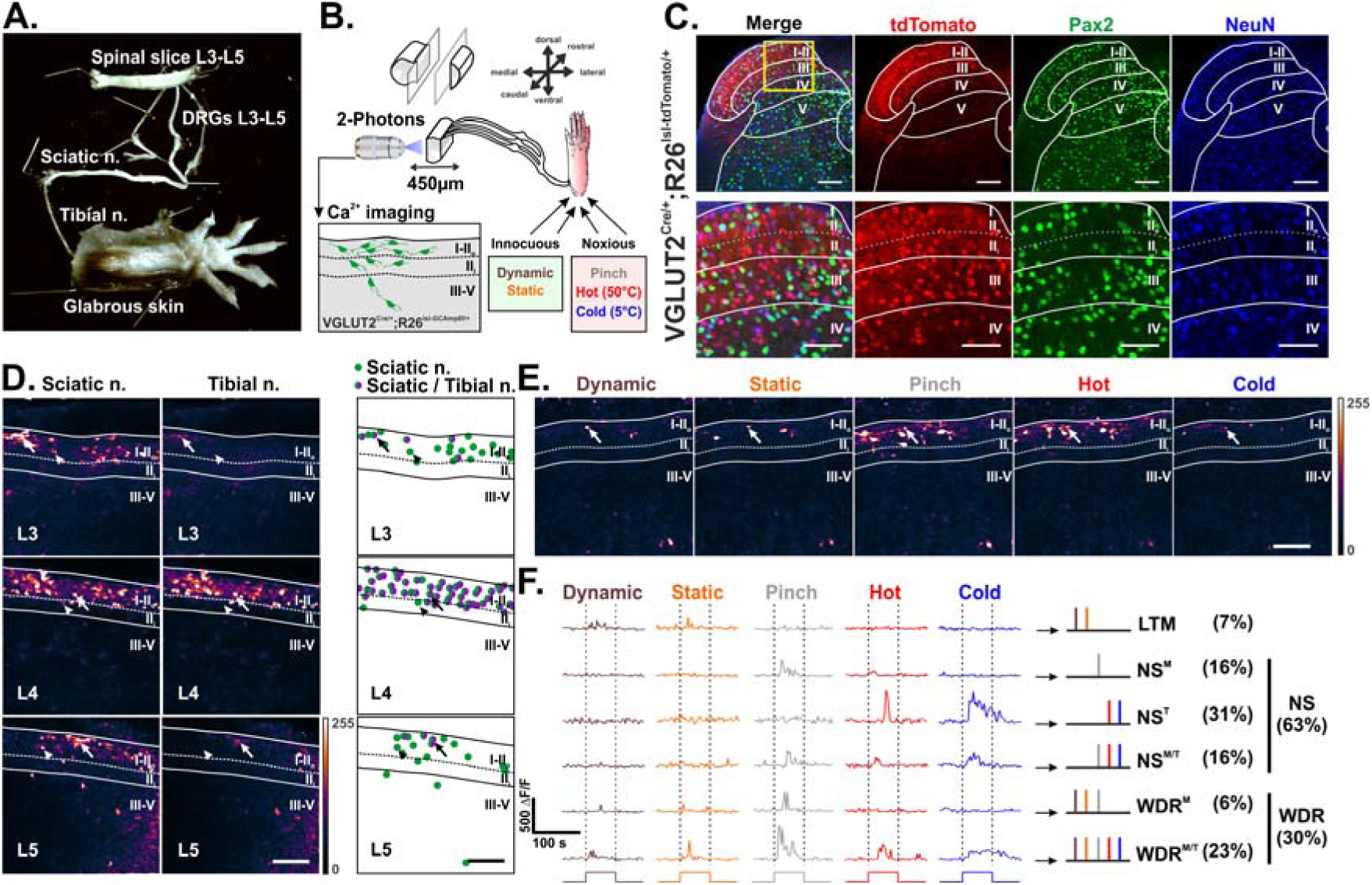
Peripheral stimulations induce Ca^2+^ transients in the DH in a new *ex vivo* spinal cord DH – skin somatosensory preparation. **(A)** Image of the *ex vivo* somatosensory preparation showing a spinal cord parasagittal slice attached to the glabrous skin of the mouse. **(B)** Schematic showing 2-photons calcium imaging performed in a DH slice of VGLUT2^cre/+^;R26^lsl-GCaMP6f/+^ mouse, following innocuous static and mechanical stimuli, or noxious mechanical, heat and cold stimuli. **(C)** Colocalization of tdTomato^+^ neurons with NeuN but not Pax2 in VGLUT2^cre/+^;R26^lsl-tdTomato/+^ mice shows that the transgene is selectively expressed in excitatory neurons. Yellow box shows location of insert. Boundaries of laminae I-II, III, IV and V are indicated by continuous lines. Borders between lamina II_o_ and II_i_ are indicated by dotted lines. Scale bars = 50 μm and 100 μm. **(D)** Representative ΔF/F images of transients evoked in the DH at the level of L3, L4 or L5 in response to suprathreshold electric stimulation of the sciatic nerve (left) or its tibial branch (right). Right panels represent cells that are activated after sciatic (green dots), or both sciatic and tibial nerve (green / magenta dots) stimulations. Arrowheads and arrows show respective examples. Boundaries of laminae I-II_o_ and III-V are indicated by continuous lines. Borders between lamina II_o_ and II_i_ are indicated by dotted lines. Look up table (LUT) of pixel intensities is indicated at the bottom right. Scale bar = 100 µm. **(E)** Representative images of Ca^2+^ transients evoked in the DH following physiological stimulations of the skin. Boundaries of laminae I-II_o_ and III-V are indicated by continuous lines. Borders between lamina II_o_ and II_i_ are indicated by dotted lines. Look up table (LUT) of pixel intensities is indicated at the bottom right. Scale bar = 100 µm. **(F)** Analysis of Ca^2+^ positive responses revealed neurons sensitive to low-threshold stimuli (LTM), to nociceptive specific stimuli (NS), or neurons with wide dynamic range responses (WDR). NS and WDR neurons are mechano- (M), thermo- (T) or mechano- and thermo- (M/T) sensitive. Each line represents transients evoked in the same neuron. Duration of stimulations are indicated beneath the traces and with dotted lines.

To validate whether VGLUT2^cre^ is strictly expressed in excitatory DH neurons, we first analyzed the expression of tdTomato in DH neurons of the reporter mouse VGLUT2^Cre/+;^R26^lsl-tdTomato/+^. Using immunofluorescence to reveal either all neurons with the neuronal marker NeuN, or only inhibitory neurons using the marker Pax2 (Peirs et al., 2020), we confirmed that the reporter is expressed in the large majority (94.7 %) of excitatory neurons from superficial to deeper laminae, but never in inhibitory neurons (Fig.1C and Table 1). It is important to note that the density of excitatory cells strongly decreased in the dorso-ventral axis, with only scattered cells in lamina V, but the ratio excitatory/inhibitory neurons remained similar across laminae (Table 1). Nevertheless, these results show that the VGLUT2^Cre^ allele captures the majority of excitatory DH neurons.

The plantar surface of the hind paw is innervated by the tibial, sural and peroneal branches of the sciatic nerve, which project to the lumbar DH through neurons with cell bodies located in L3 to L5 DRGs in mice (Rigaud et al., 2008). To assess the functional connectivity of the hind paw to the DH in this somatosensory preparation, we performed Ca^2+^ imaging of L3-L5 DRGs and the related lumbar spinal cord, while stimulating electrically the sciatic nerve, or only its tibial branch, which innervates most of the glabrous pad. To address whether conditional expression of GCamp6f in VGLUT2+ cells could be reliably used to report gross activity of DRGs neurons, we first analyzed which peripheral sensory populations expressed the transgene in the VGLUT2^Cre/+;^ R26^lsl-tdTomato/+^ reporter mouse. Similar to previous observations (Scherrer et al., 2010), we found that the reporter is expressed in the main populations of sensory neurons, *i.e.* in 72 ± 7 % of myelinated neurons expressing the neurofilament 200 (NF200), and in 87 ± 6 % and 96 ± 2 % of unmyelinated peptidergic neurons positive for the calcitonin gene-related peptide (CGRP), or non peptidergic neurons positive for the Isolectine B4 (IB4), respectively (Fig. S1A). We next analyzed Ca^2+^ transients evoked by suprathreshold electric stimulations of the sciatic nerve in the DRGs and lumbar DH of VGLUT2^Cre/+;^ R26^lsl-GCamp6f/+^ mice. In line with previous anatomical and functional studies, we found numerous activated cells after stimulations of the sciatic nerve at the level of L3-L5 in both peripheral and central regions (Figs. 1D and S1B). In contrast, the vast majority of neurons activated by stimulations of the tibial nerve were located in the L4 DRG and the respective spinal segment, with only scattered cells in adjacent regions (Fig. 1D). As expected from our observations with the reporter mouse (Fig. 1C), the cell density was strongly higher in the superficial laminae compared to the deep DH, especially in the region located above neurons of the inner lamina II (II_i_) that express the gamma isoform of protein kinase C (PKCγ) (Figs. 1D and S1C).

Finally, we recorded Ca^2+^ transients in L4 DH neurons evoked by physiological stimulations of the hind paw of VGLUT2^Cre/+;^ R26^lsl-GCamp6f/+^ mice. The skin was stimulated using innocuous dynamic and static mechanical stimuli, and noxious mechanical and thermal heat and cold stimuli (Fig. S1D). We classified neurons as low-threshold mechanosensitive (LTM) if they responded only to innocuous stimuli, noxious specific (NS) if they responded to only noxious stimuli, and wide dynamic range (WDR) responders, also known as convergent or multi-receptive neurons, for those responding to at least one noxious and one innocuous simulation (Figs. 1E-F), as described in similar studies (Zhang et al., 2013). Under these conditions, 69% of all responding neurons were mechanosensitive, consistent with a recent study that analyzed the sensory modality of superficial DH neurons (Warwick et al., 2022). Importantly, we found that 63% of responsive neurons were NS, 30% were WDR and 7% were LTM (Fig.1F), which was nearly identical to previous observations from the superficial laminae using extracellular electrophysiology *in vivo* (Zhang et al., 2013).

Altogether, these data show that this new adapted somatosensory preparation is highly reliable to address spinal cord excitatory circuits activity across the entire DH, following physiological stimulations of the skin.

### Aδ and C-fiber mediated neuronal activity predominates in superficial DH neurons under physiological conditions

Noxious and innocuous sensory information are conveyed to DH neurons through various peripheral sensory populations with specific organ endings (Peirs & Seal, 2016). These sensory populations display different activation thresholds and levels of myelination, and are typically classified as Aβ low-threshold (LT) sensory neurons for highly myelinated neurons, Aδ high-threshold (HT) sensory neurons or Aδ-LT for thinly myelinated neurons, and C-HT or C-LT for unmyelinated neurons (Fig. 2A). Interestingly, these populations of sensory neurons can be activated by electric stimulations of the dorsal roots, with ranges of intensities that can recruit preferentially only Aβ, Aβ + Aδ, or Aβ + Aδ + C fibers, respectively (Baba et al., 1999). We thus assessed which DH neurons were activated by these distinct populations of afferents in VGLUT2^cre/+;^R26^lsl-GCaMP6f/+^ mice (Fig. 2B).

**Figure 2.**
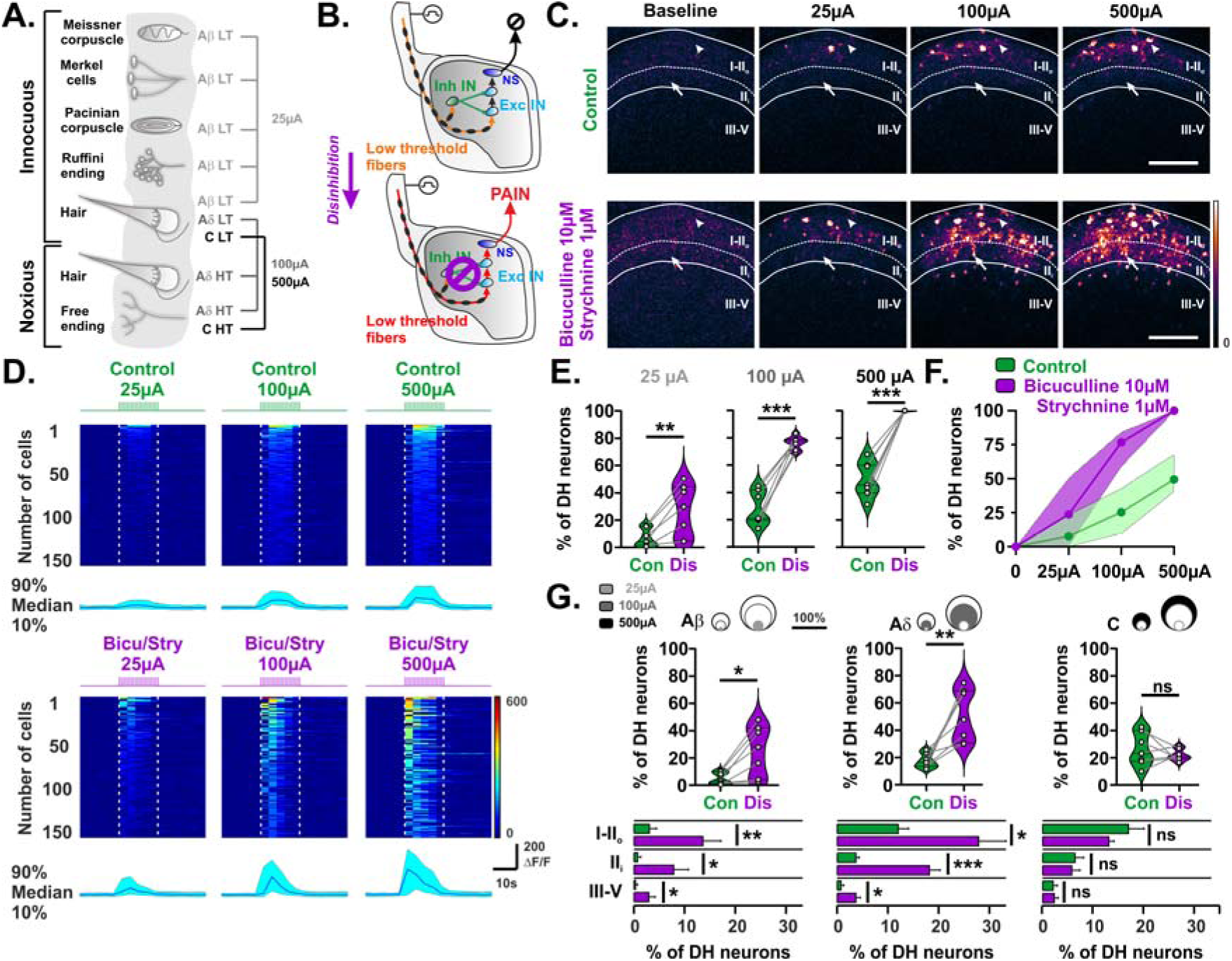
Spinal disinhibition sensitizes DH neurons to A-fiber inputs from lamina V to I. **(A)** Schematic representation of peripheral sensory neurons innervating the skin, and their sensitivity to electric stimulations at 25, 100 and 500 µA. **(B)** Schematic showing that in control conditions (above), low threshold sensory neurons (orange) do not activate excitatory interneurons (Exc IN) or nociceptive specific neurons (NS) due to local inhibitory neurons (IN). Pharmacological disinhibition (below) allows low threshold sensory neurons (red) to activate NS neurons through polysynaptic Exc IN, thus inducing pain. **(C)** Representative images of Ca^2+^ transients evoked in the DH following electric stimulations of the dorsal root at 25, 100 and 500 µA to activate Aβ, Aδ and C-fibers respectively, in control conditions (top) and after spinal disinhibition (bottom). Arrowheads and arrows show examples of cells activated by stimulations of A-fibers in control conditions, or only after disinhibition, respectively. Boundaries of laminae I-II_o_ and III-V are indicated by continuous lines. Borders between lamina II_o_ and II_i_ are indicated by dotted lines. Look up table (LUT) of pixel intensities is indicated at the bottom right. Scale bar = 100 µm. **(D)** Heatmaps showing ΔF/F Ca^2+^ transients of 150 representative neurons during dorsal root stimulations at 25, 100 and 500 µA in control conditions (top) and after spinal disinhibition (bottom). Each line represents the same cell during each condition. Median, 10 and 90 percentiles of the amplitude of these transients are shown below each heatmap. Duration of stimulations are indicated with dotted lines. LUT of the heatmaps is indicated at the bottom right. Bicu; Bicuculline. Stry; Strychnine. **(E)** Absolute or **(F)** Cumulative percentage of DH neurons responding to electric stimulations of the dorsal root at 25, 100 and 500 µA in control conditions (Con) or after spinal disinhibition (Dis). n=7 mice. **p ≤ 0.01, ***p ≤ 0.001, paired t tests. **(G)** Percentage of DH neurons with Aβ- (± Aδ ± C), Aδ-(± C) or C-fiber inputs in control conditions (Con) or after spinal disinhibition (Dis). Pie charts above represent the proportion of these neurons that begin to respond at 25 µA (Aβ threshold, light grey), then 100 µA (Aδ threshold, dark grey) and finally 500 µA (C threshold, black). Distributions of these neurons across laminae I-II_o_, II_i_ or III-V are indicated below each graph. n=7 mice. ns= not significant, *p < 0.05, **p ≤ 0.01, ***p ≤ 0.001, paired t tests and matched-paired Wilcoxon tests.

In control conditions, very few neurons (6.9% ± 2.5 of all DH neurons) exhibited Ca^2+^ transients after stimulation of Aβ afferents with 25µA, and those were preferentially located in laminae I and the outer lamina II (II_o_) (Figs. 2C-E). It is also important to note that neuronal activity was detected nearly exclusively during the stimulation period, indicating low spontaneous activity or post-discharge in DH neurons in these conditions (Fig. 2D). Increasing the intensity of stimulations up to 100 µA in order to additionally recruit Aδ afferents, or up to 500 µA to also recruit C afferents, elicited transients in 28.1% ± 4.7 and 49.7% ± 4.9 of DH neurons respectively, which were again located preferentially in the most superficial laminae (Figs. 2C-E). Notably, increasing the intensity of stimulations affected not only transient amplitudes (Fig. S2A), but also the number of responding neurons (Fig. 2F), suggesting that Aβ-, Aδ- and C-fibers recruit convergent but also distinct DH circuits in control conditions. While we could not distinguish between cells that receive only one, or multiple types of afferents, because increasing the intensity of electric stimulations will only recruit additional cells, we nonetheless separated neurons that received only C afferents, and those that receive Aδ (± Aβ) or Aβ (± Aδ ± C) fibers inputs. Using this segmentation, we found that very few neurons were activated by Aβ stimulations in control conditions (5.0% ± 1.7 of DH excitatory neurons), and those were located mostly in laminae I-II_o_ (Fig. 2G). In contrast, neurons receiving Aδ- and C-fibers inputs represented 17.4% ± 2.0 and 26.2% ± 4.6 of DH excitatory neurons respectively, with about 2/3 located in lamina I-II_o_, and 1/3 located mostly in lamina II_i_ (Fig. 2G).

Altogether, these results indicate that, in control conditions, excitatory DH neurons are mostly activated in superficial laminae by stimuli of high intensities, which likely recruit preferentially Aδ-HT and C-HT sensory neurons, consistent with previous functional analysis of DH neurons using activity markers (Ji et al., 1999), Ca^2+^ imaging (Sullivan and Sdrulla, 2022) or multiarray electrophysiological recordings (Greenspon et al., 2019).

### Spinal disinhibition unmasks Aβ- and Aδ-fiber mediated neuronal activity in both superficial and deep DH neurons

Dorsal roots were next stimulated after spinal disinhibition, a well-characterized model of touch-evoked mechanical allodynia (Baba et al., 2003; Miraucourt et al., 2007; Torsney and MacDermott, 2006), and a hallmark of neuropathy (Gradwell et al., 2019; Latremoliere and Woolf, 2009). We performed spinal disinhibition using bath application of Bicuculine (10 µM) and Strychnine (1 µM) to inhibit gamma-aminobutyric acid type A receptors (GABA_A_) and glycinergic receptors, respectively.

Interestingly, in the absence of peripheral stimuli, no spontaneous activity was observed 30 minutes after drugs application, suggesting that there was relatively low segmental tonic inhibition of excitatory DH neurons under control conditions (Figs. 2C-D), as previously reported *in vivo* (Lee et al., 2019). In contrast, under spinal disinhibition, the number of responding neurons was dramatically increased following stimulations of the dorsal root with 25, 100 and 500 µA respectively (Figs. 2C-E). The proportion of responding neurons was also affected, increasing the number of cells responding to 25 µA and decreasing those responding to only 500 µA (Fig. S2B), with no significant change in the amplitude of their Ca^2+^ transients (Fig. S2A). Interestingly, when we performed again a segmentation of responders based on their afferent inputs, we found that spinal disinhibition significantly increased the number of neurons receiving Aβ- and Aδ-, but not C-fiber inputs (Fig. 2G). Most importantly, Aβ- and Aδ-fiber stimulations significantly recruited more neurons in superficial laminae I-II_o_, but also in lamina II_i_ and in deeper laminae, after spinal disinhibition (Fig. 2G). Unexpectedly, we found that among all responding neurons after disinhibition, 50.3% ± 4.9 were previously silent neurons that were not responding to suprathreshold electric stimulations during control conditions. These newly recruited neurons mainly received Aδ-fibers (56.8%) and to a less extent C- (28.8%) or Aβ- (14.1%) fibers. Those were equally distributed between laminae I-II_o_ and LII_i_ (46.3% and 41.4% of DH neurons respectively) while the remaining 11.6% were located in the deep DH.

Altogether, these data indicate that spinal disinhibition unmasks neurons responding to Aβ- and Aδ-fibers across all layers of the DH, with about half coming from a pool of previously silent neurons.

### Spinal disinhibition sensitizes superficial DH neurons to innocuous and noxious peripheral stimuli

Electrophysiological, transcriptomic and functional analysis of sensory neurons have shown that A-fibers can relay a variety of innocuous, as well as noxious, mechanical and thermal information (Wang et al., 2023). Since we showed that spinal disinhibition sensitizes DH neurons predominantly to A-fiber inputs (Fig. 2), we further characterized more precisely which sensory modality could be affected by loss of spinal inhibition. As in Fig.1, we stimulated the hind paw of VGLUT2^cre/+^;R26^lsl-GCaMP6f/+^ mice using innocuous dynamic and static mechanical stimuli, and noxious mechanical and thermal heat and cold stimuli, and recorded DH Ca^2+^ transients.

In control conditions, only 33.4% ± 3.3 of all DH excitatory neurons responded to at least one physiological peripheral stimulus, suggesting that most of the DH is either under strong inhibitory control, or tuned for other sensory modalities not tested in our experimental paradigm (Figs. 3A-C). Similar to what we observed with electric stimulations of the sciatic nerve in control conditions, responding neurons were predominantly found in the most superficial laminae (Figs. 3A-C). Given that the density of excitatory DH neurons decreases through the dorso-ventral axis (Table 1), we further normalized the percentage of responding cells by lamina. Interestingly, such segmentation revealed that a similar proportion of cells within each lamina participated in sensory processing, with 37.7% ± 4.2 of lamina I-II_o_, 23.8% ± 5.3 of lamina II_i_ and 35.5%± 17.9 of lamina III-V responding to at least one stimulus.

**Figure 3.**
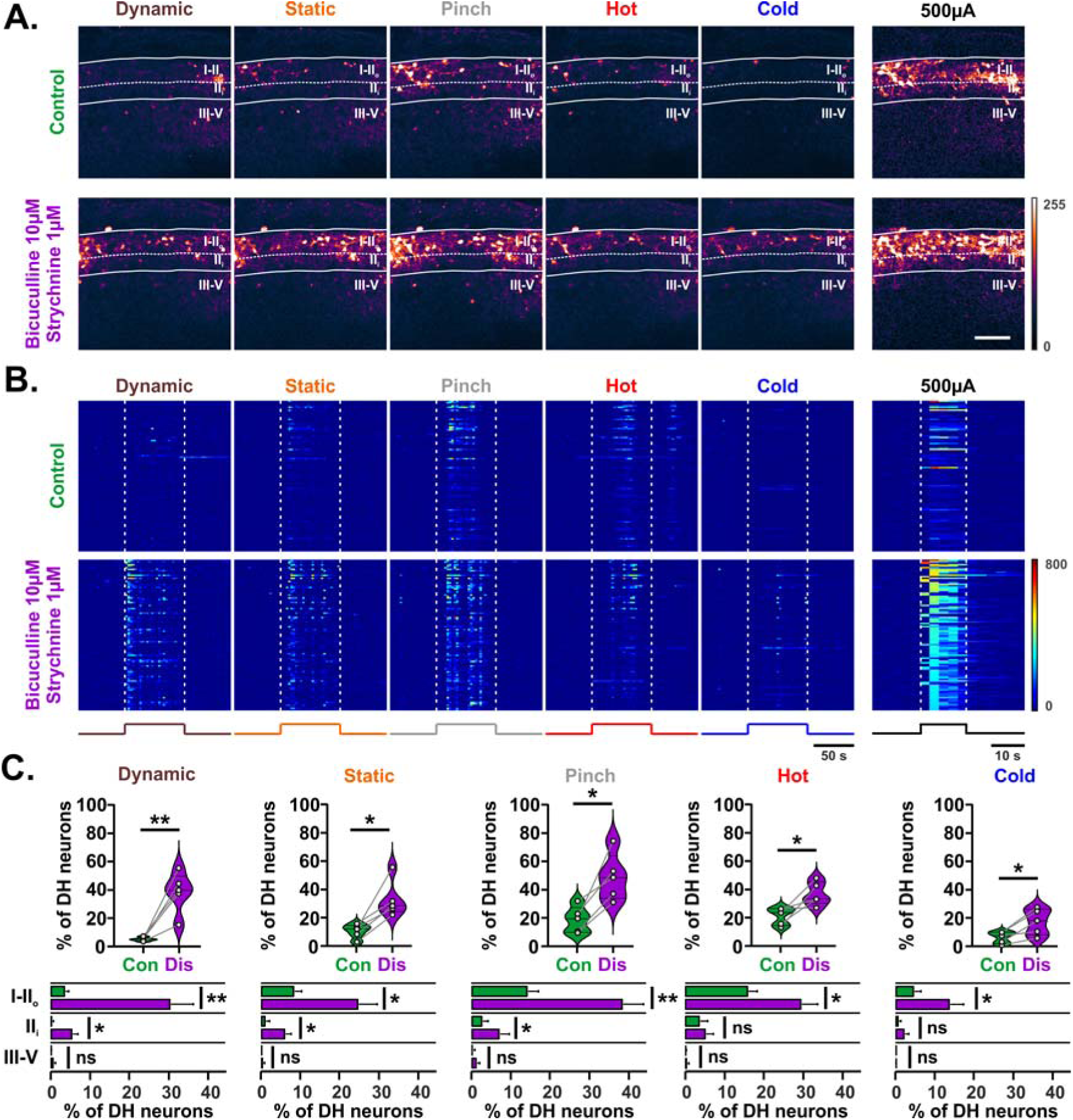
Spinal disinhibition leads to hyperactivity of DH neurons following mechanical stimuli in lamina I-II_i_ and thermal stimuli in laminae I-II_o_. **(A)** Representative images of Ca^2+^ transients evoked in the DH following physiological stimulations of the skin, or suprathreshold electric stimulation of the tibial nerve, in control conditions (above) and after spinal disinhibition (bottom). Boundaries of laminae I-II_o_ and III-V are indicated by continuous lines. Borders between lamina II_o_ and II_i_ are indicated by dotted lines. Look up table (LUT) of pixel intensities is indicated at the bottom right. Scale bar = 100µm. **(B)** Heatmaps showing ΔF/F Ca^2+^ transients of 100 representative neurons physiological stimulations of the skin in control conditions (top) and after spinal disinhibition (bottom). Each line represents the same cell during each condition. Median, 10 and 90 percentiles of the amplitude of these transients are shown below each heatmap. Duration of stimulations are indicated with dotted lines. LUT of the heatmaps is indicated at the bottom right. **(C)** Percentage of DH neurons responding to physiological stimulations of the skin in control conditions (Con) or after spinal disinhibition (Dis). Distributions of these neurons across laminae I-II_o_, II_i_ or III-V are indicated below each graph. n=5 mice. ns= not significant, **p < 0.01, *p < 0.05, paired t tests and matched-paired Wilcoxon tests.

We next applied Bicuculine and Strychnine, as described above. Similar to what we saw with dorsal root electric stimulations (Fig. 2), responding neurons were far more numerous after disinhibition (Figs 3A-C), with now up to 70.6% ± 3.1 of all DH excitatory neurons responding to at least one peripheral stimulation. Such increase in neuronal activity was observed regardless of the modality tested (Fig 3C), suggesting that there is virtually no sensory input that lacks inhibitory controls at the level of the DH. Importantly, 51.6% ± 6.9 of transients recorded after disinhibition came from previously silent neurons. Analysis of the amplitude of individual Ca^2+^ transients revealed only mild changes after spinal disinhibition (Fig. S3), suggesting that spinal disinhibition preferentially regulates the number of neurons responding to peripheral stimuli, rather than the Ca^2+^ activity within an already responding cell. Similar to what we observed under control conditions, a relatively homogenous contribution from each lamina was recorded following stimulations of the skin, with 75.9% ± 2.7 of lamina I-II_o_, 63.5% ± 8.7 of lamina II_i_ and 63.7% ± 10.9 of lamina III-V neurons responding to at least one stimulus. Again, the highest activity was recorded in lamina I-II_o_ after disinhibition (Figs. 3C). We also observed a significant increase in the activity of excitatory DH neurons of lamina II_i_, particulary following innocuous mechanical stimuli, with a 9.3 ± 3.4 fold increase following dynamic, and a 6.3 ± 1.6 fold increase following static stimuli (Figs. 3C). Of note, no significant change in the number of responsive cells were observed in the deepest DH, no matter what stimulus was applied, suggesting that less plasticity of sensory circuits occurs in this area (Fig. 3C).

Together, these data indicate that spinal disinhibition sensitized superficial DH excitatory neurons to innocuous and noxious stimuli, with cells in lamina II_i_ being newly activated mainly by mechanical sensory inputs.

### Spinal disinhibition increases the number of WDR neurons and decreases the number of silent neurons

We showed that spinal disinhibition led to DH hyperactivity following peripheral stimulations. Such hypersensitivity could result from the recruitment of previously silent neurons, or gain-of-function of responding cells. We thus analyzed each individual neuronal response to the five peripheral stimulations, resulting in 32 possible patterns, ranging from no response at all, to response following each stimulus. We then compared these 32 patterns before and after disinhibition, resulting in 32 x 32 = 1024 possible transitions. Given this high number of combinations and to achieve a meaningful statistical power, neuronal responses were analyzed together from 5 animals, totaling 1106 DH excitatory neurons.

The sensory modalities of more than half of excitatory neurons were altered by spinal disinhibition, while 44% responded to the same stimuli before and after drugs application (Figs. 4A and S4). 35% of neurons became WDR neurons, with about 2/3 coming from a pool of previously silent neurons (Fig. 4A). In contrast, only 14 and 6% of neurons became NS and LTM, respectively. Such plasticity in sensory processing was not homogenous between DH laminae. While silent and NS neurons were clearly the most numerous within all laminae in control conditions, WDR neurons were far more predominant in laminae I-II_o_ after disinhibition, making up to 67% of all responding neurons in this area (Figs. 4B-C). WDR neurons also represented the largest class of responding cells in deeper laminae after spinal disinhibition, making about half of all responding neurons, but those were still outnumbered by silent neurons even in the absence of inhibition (Fig. 4B). As a result from the increase in the proportion of WDR neurons following disinhibition, NS neurons represented only 27% of all DH responding neurons after disinhibition, compared to 63% in control conditions (Fig S5). Of note, the proportion of LTM neurons was nearly identical in laminae I-II_o_, or III-V before and after disinhibition, but increased by more than 3 times in lamina II_i_ (Fig. 4C), suggesting that lamina II_i_ is particularly sensitive to hyperexcitability following stimulations of innocuous mechanical inputs.

**Figure 4.**
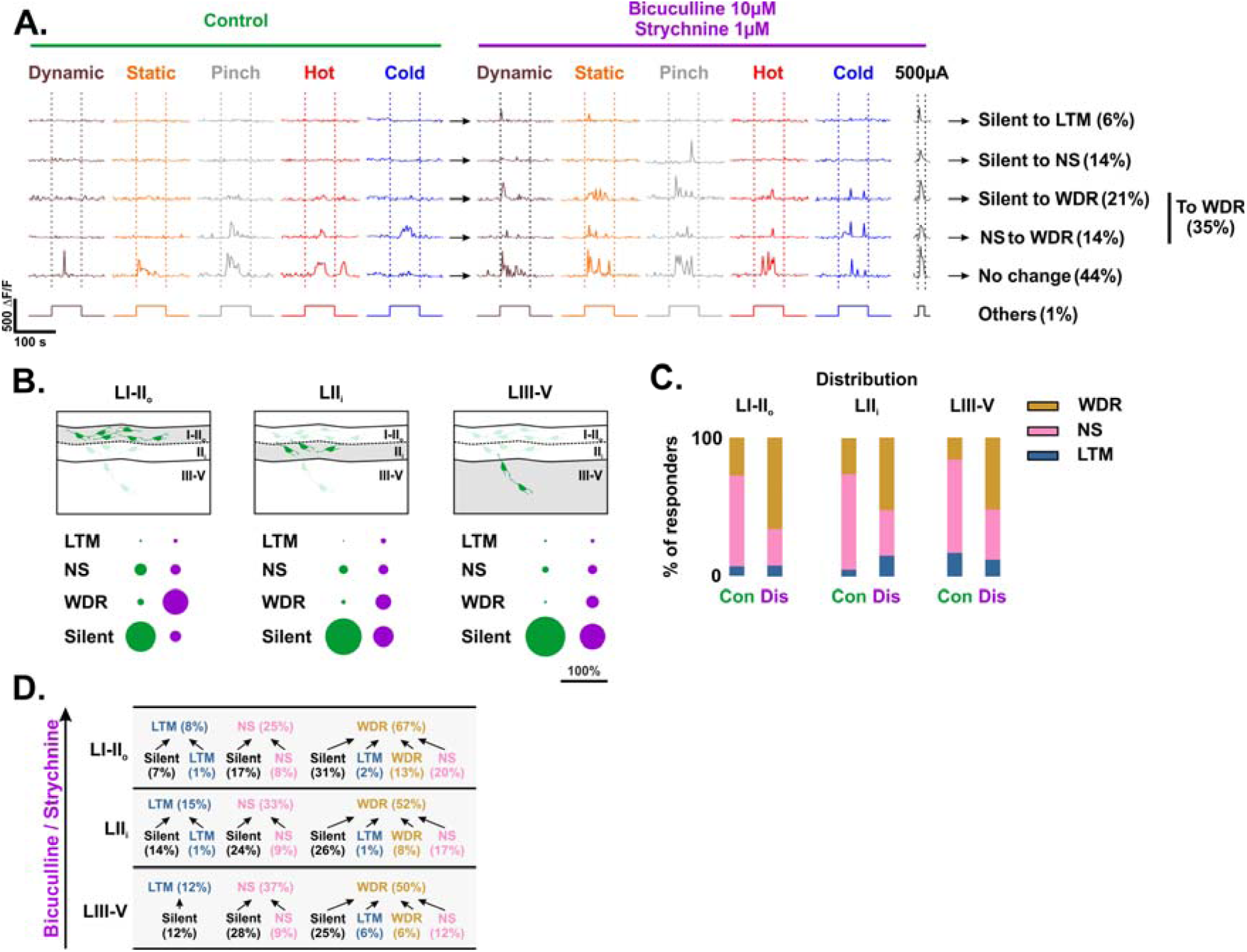
Spinal disinhibition unmasks a large number of previously silent neurons and turns most DH neurons into polymodal cells. **(A)** Representative analysis of Ca^2+^ transients revealing the major changes in sensory processing by DH neurons observed after spinal inhibition. These include previously silent neurons that began to respond to low-threshold mechanical stimuli (LTM), to nociceptive specific stimuli (NS), or neurons with wide dynamic range responses (WDR) after spinal disinhibition. Some NS neurons became WDR after spinal disinhibition, while the remaining responded similarly as in control conditions. Each line represents transients evoked in the same neuron. Duration of stimulations are indicated beneath the traces and with dotted lines. **(B)** Proportions of LTM, NS, WDR and silent neurons in control conditions and after spinal disinhibition within laminae I-II_o_, II_i_ or III-V. **(C)** Distribution of responding neurons in control conditions (Con) and after spinal disinhibition (Dis) within laminae I-II_o_, II_i_ or III-V. **(D)** Framework of the DH network showing the major transformations in sensory processing observed after spinal disinhibition within laminae I-II_o_, II_i_ or III-V. Arrows point toward the proportions of LTM, NS, WDR neurons observed after spinal disinhibition, and originate from what they were in control conditions.

Altogether, these data indicate that loss of spinal inhibition preferentially reduces the number of silent neurons and increases the number of neurons responding to innocuous mechanical input. Those, which include the LTM and WDR cells, now account for up to 73% of DH responding neurons, compared to 37% in control conditions.

### Spinal inhibition primarily turns silent neurons into WDR neurons

Given that most DH excitatory neurons have the ability to receive numerous sensory information (Fig. S4), their sensory modality seems to be finely tuned by inhibitory interneurons that act on these inputs. We thus next aimed at building a framework of these inhibitory controls onto the various populations of DH neurons, to map how sensory inputs are processed in control conditions and after loss of spinal inhibition. To perform this analysis, we isolated the main LTM, NS, WDR and silent populations of neurons that were identified in control conditions, and analyzed what each of them became after disinhibition (Fig S6A). We then compared the major populations of neurons identified after disinhibition, to what they were in control conditions (Fig S6B). The major transitions were then assessed individually to reveal which of their sensory inputs is controlled by spinal inhibition (Fig. S7).

We identified five main classes of neurons. Three were found in all laminae, and included two populations of silent neurons that respectively turned into NS or WDR neurons after loss of inhibition (Fig. S7, Silent->NS and Silent->WDR), and a third population of NS neurons that became WDR after loss of inhibition (Fig. S7, NS->WDR). The fourth population was predominantly found from lamina II_i_ to V and included silent neurons that became LTM after loss of inhibition (Fig. S7, Silent->LTM). The last population was mostly found in laminae I-II_o_ and included neurons that were WDR in all conditions (Fig. S7, WDR).

Consistent with the strong decrease in the number of silent neurons after spinal disinhibition, we found that more than half of the activity of DH neurons came from previously silent neurons under these conditions. Silent->WDR were predominant in laminae I-II_o_, representing 31% of responding neurons in this area (Fig. 4D). Those represented about 25% of responding neurons in deeper laminae, neighboring an equal proportion of Silent->NS neurons. Interestingly, there was an increasing number of NS->WDR neurons from the deep to the most superficial laminae, but this gradient was inverted for Silent->LTM neurons. This latter observation coincides well with previous studies suggesting that spinal disinhibition leads to the recruitment of previously silent neurons from the deep DH by innocuous inputs, that would feed onto the nociceptive circuit of the superficial laminae (Miraucourt et al., 2007; Torsney and MacDermott, 2006). The data however also shows that this mechanism represents only a small fraction of the sensory plasticity induced by spinal disinhibition (Fig. 4D). We thus conclude that spinal disinhibition primarily leads to four major changes in sensory processing within DH neurons: i) Silent->WDR neurons within all laminae of the DH, ii) NS->WDR in the superficial DH, and iii) silent->LTM and iv) Silent->NS in the deep DH.

### Nerve injury enhances Aβ-, but not Aδ- or C-fiber mediated neuronal activity, selectively in superficial DH neurons

An extensive body of work suggests that central sensitization of the DH, a hallmark of neuropathy, is closely related to the alteration of DH inhibitory controls (Zeilhofer and Ganley, 2019). We next assessed whether peripheral nerve injury would recapitulate the main features that we observed after spinal disinhibition, i.e. facilitation of A-fiber mediated activation of lamina I-V neurons and sensitization of superficial DH neurons to innocuous mechanical, and noxious mechanical and thermal stimuli. We used the cuff model of persistent neuropathic pain (Fig. 5A), which tightly mimics the symptoms, comorbidities and pharmacological therapeutic sensitivity of neuropathic chronic pain (Kremer et al., 2021).

**Figure 5.**
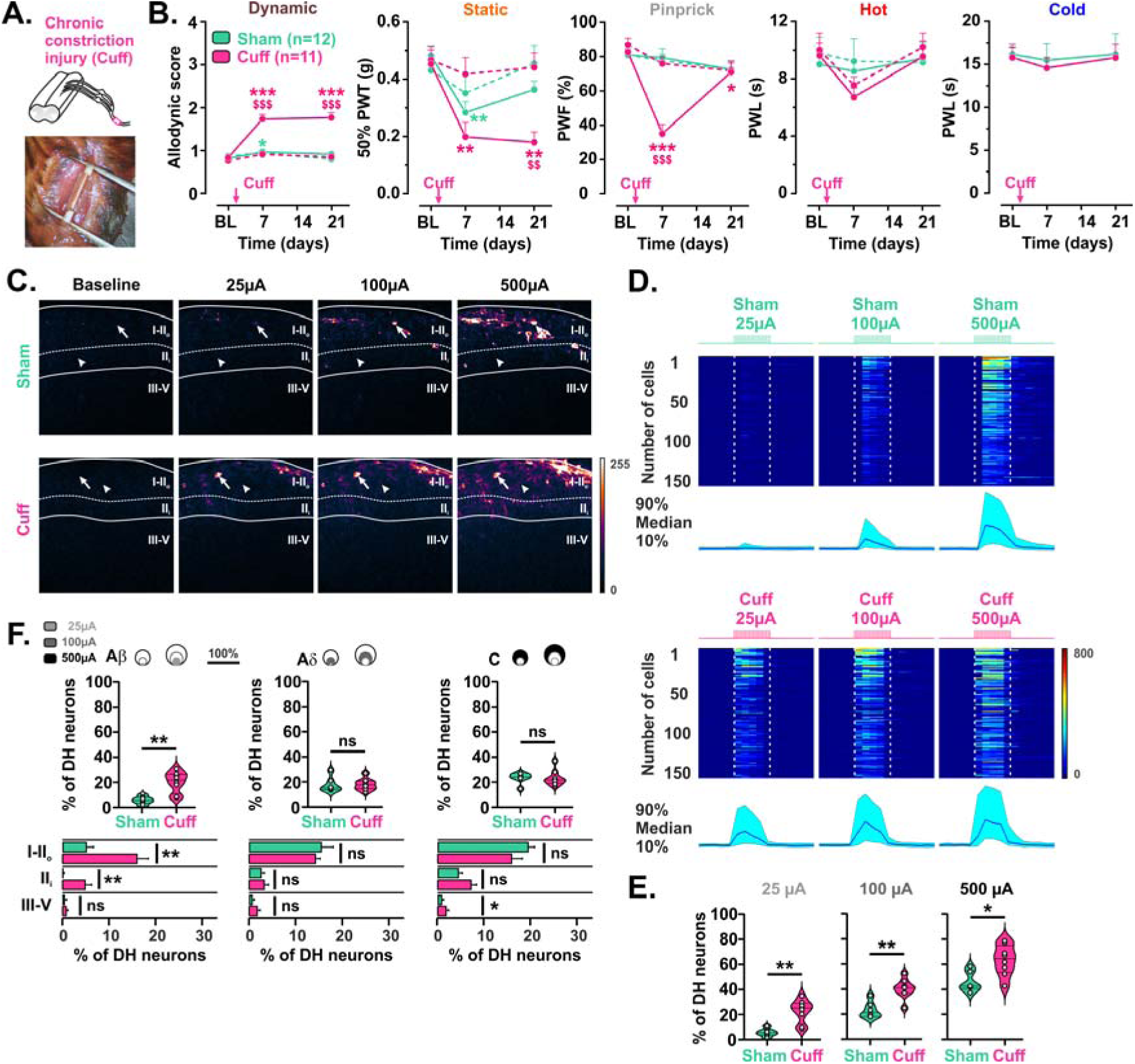
Nerve injury sensitizes DH neurons to Aβ-fiber inputs from lamina II_i_ to I. **(A)** Schematic and representative photograph of the Cuff model showing a permanent polyethylene tubing placed around the sciatic nerve. **(B)** Behavioral analysis of sensitivity to innocuous mechanical dynamic and static stimuli, and to noxious mechanical and thermal stimuli, in Sham vs Cuff mice, in control conditions and up to 3 weeks after surgery. n=12 or n=6 per group for mechanical or thermal tests, respectively. Plain and dotted line represent the ipsilateral and contralateral sides, respectively. *p < 0.05, **p < 0.01 and ***p < 0.001 for comparisons within animal groups to baseline values. ^$$^p < 0.01 and ^$$$^p < 0.001 for comparisons between Sham and Cuff values on the ipsilateral side across time. 2-way ANOVA with repeated measures and Dunnett’s (*) or Tuckey (^$^) post hoc tests. **(C)** Representative images of Ca^2+^ transients evoked in the DH following electric stimulations of the dorsal root at 25, 100 and 500 µA to activate Aβ, Aδ and C-fibers respectively, in Sham operated mouse (top) and Cuff mouse (bottom). Arrows show examples of cells activated by stimulations of Aδ-fibers in Sham or Aβ-fibers in Cuff animals. Arrowheads show examples of cells activated in Sham or Cuff animals only after an additional pharmacological spinal disinhibition. Boundaries of laminae I-II_o_ and III-V are indicated by continuous lines. Borders between lamina II_o_ and II_i_ are indicated by dotted lines. Look up table (LUT) of pixel intensities is indicated at the bottom right. Scale bar = 100 µm. **(D)** Heatmaps showing ΔF/F Ca^2+^ transients of 150 representative neurons during dorsal root stimulations at 25, 100 and 500 µA in Sham (top) vs Cuff (bottom) animals, and at 500 µA after an additional pharmacological spinal disinhibition. Each line represents the same cell during each condition. Median, 10 and 90 percentiles of the amplitude of these transients are shown below each heatmap. Duration of stimulations are indicated with dotted lines. LUT of the heatmaps is indicated at the bottom right. **(E)** Absolute percentage of DH neurons responding to electric stimulations of the dorsal root at 25, 100 and 500 µA in Sham vs Cuff animals. Sham n=6, Cuff n=8 mice. *p < 0.05, **p ≤ 0.01, unpaired t tests. **(F)** Percentage of DH neurons with Aβ- (± Aδ ± C), Aδ- (± C) or C-fiber inputs in Sham vs Cuff animals. Pie charts above represent the proportion of these neurons that begin to respond at 25 µA (Aβ threshold, light grey), then 100 µA (Aδ threshold, dark grey) and finally 500 µA (C threshold, black). Distributions of these neurons across laminae I-II_o_, II_i_ or III-V are indicated below each graph. Sham n=6, Cuff n=8 mice. ns= not significant, *p < 0.05, **p ≤ 0.01, unpaired t tests and Mann-Whitney tests.

We first assessed mice sensitivity to innocuous static and dynamic mechanical stimuli, and to noxious mechanical and thermal heat and cold stimuli using behavioral tests. As previously reported using the cuff model (Yalcin et al., 2014), animals developed an ipsilateral static mechanical allodynia as early as 1 week following the surgery, which lasted for at least three weeks (Fig. 5B). In addition, mice developed a long-lasting dynamic mechanical allodynia (Fig. 5B), similar to what was reported using other neuropathic models (Duan et al., 2014). We also observed a transient but pronounced mechanical hypoesthesia, a highly prevalent symptom of postoperative pain (Borsook et al., 2013). In contrast, mice did not show any long-lasting difference in noxious thermal sensitivity compared to sham animals using paw immersion tests (Fig. 5B) or noxious thermal plates (Fig. S8A), as previously reported with the cuff model (Benbouzid et al., 2008; Nascimento et al., 2015) but see (Bailey and Ribeiro-da-Silva, 2006; Cardenas et al., 2020).

Next, we analyzed how Ca^2+^ transients evoked by peripheral stimulations were affected after nerve injury in DH excitatory neurons, using VGLUT2^cre/+;^R26^lsl-GCaMP6f/+^ mice. As described above, we first performed electric stimulations of L4-5 dorsal roots. As in naïve animals (Fig. 2), very few neurons were activated in the sham group following stimulation with 25µA, and about 20 to 50% of all DH neurons were activated by 100 and 500 µA, respectively (Figs. 5C-F). Again, transients were observed mostly in the most superficial laminae I and II_o_ under these conditions (Fig. 5C). In neuropathic animals, responding neurons were much more numerous, lighting up to 22.6% ± 3.3, 40.0% ± 2.8 and 63.1% ± 4.3 of all DH neurons following stimulations with 25, 100 and 500 µA, respectively (Fig. 5E). Interestingly, when we segmented neurons to reveal those receiving Aβ- and/or Aδ- and/or C-fibers inputs, we found an increase in responding neurons only for those with Aβ-fiber inputs in neuropathic animal compared to shams (Fig. 5F), consistent with previous reports in rats (Schoffnegger et al., 2008). Those were located in laminae I-II_o_ and lamina II_i_ but not in deeper laminae (Fig. 5H).

Together, these data show that nerve injury leads to mechanical hypersensitivity associated with an increase in neuronal activity from lamina II_i_ to I.

### Nerve injury sensitizes DH neurons selectively to innocuous mechanical stimuli in the superficial DH

We next stimulated the ipsilateral hind paw of VGLUT2^cre/+^;R26^lsl-GCaMP6f/+^ neuropathic mice using the five physiological stimuli described earlier, and recorded DH Ca^2+^ transients. As in naïve animals (Fig. 2), DH neurons of sham mice were preferentially sensitive to noxious stimuli, and those were again mostly found in the most superficial laminae I-II_o_ (Fig 6A). Proportions of LTM, NS, WDR and silent neurons were also comparable in sham compared to naive animals, with 64% of all DH neurons being silent, followed by NS (24%), WDR (10%) and very few LTM (2%) neurons (Fig. S8B). In contrast, 59% of all DH neurons responded to at least one peripheral stimulus after nerve injury. Interestingly, as observed after pharmacological DH disinhibition (Fig. 4), a similar proportion (≈40%) of responding neurons were recorded within each lamina following peripheral stimuli in neuropathic animals (Fig. 6B), suggesting that spinal inhibition was reduced homogeneously within each lamina after nerve injury. In line with what we observed with behavior analysis, we found a significant increase in the total number of responding neurons after nerve injury following innocuous mechanical dynamic (+413%) and static (+155%), but also to a less extend noxious mechanical and heat stimuli (Fig. 6A). Importantly, mechanical hyperactivity was significantly observed only in lamina I-II_i_, but not in the deepest DH compared to sham animals (Fig. 6A).

**Figure 6.**
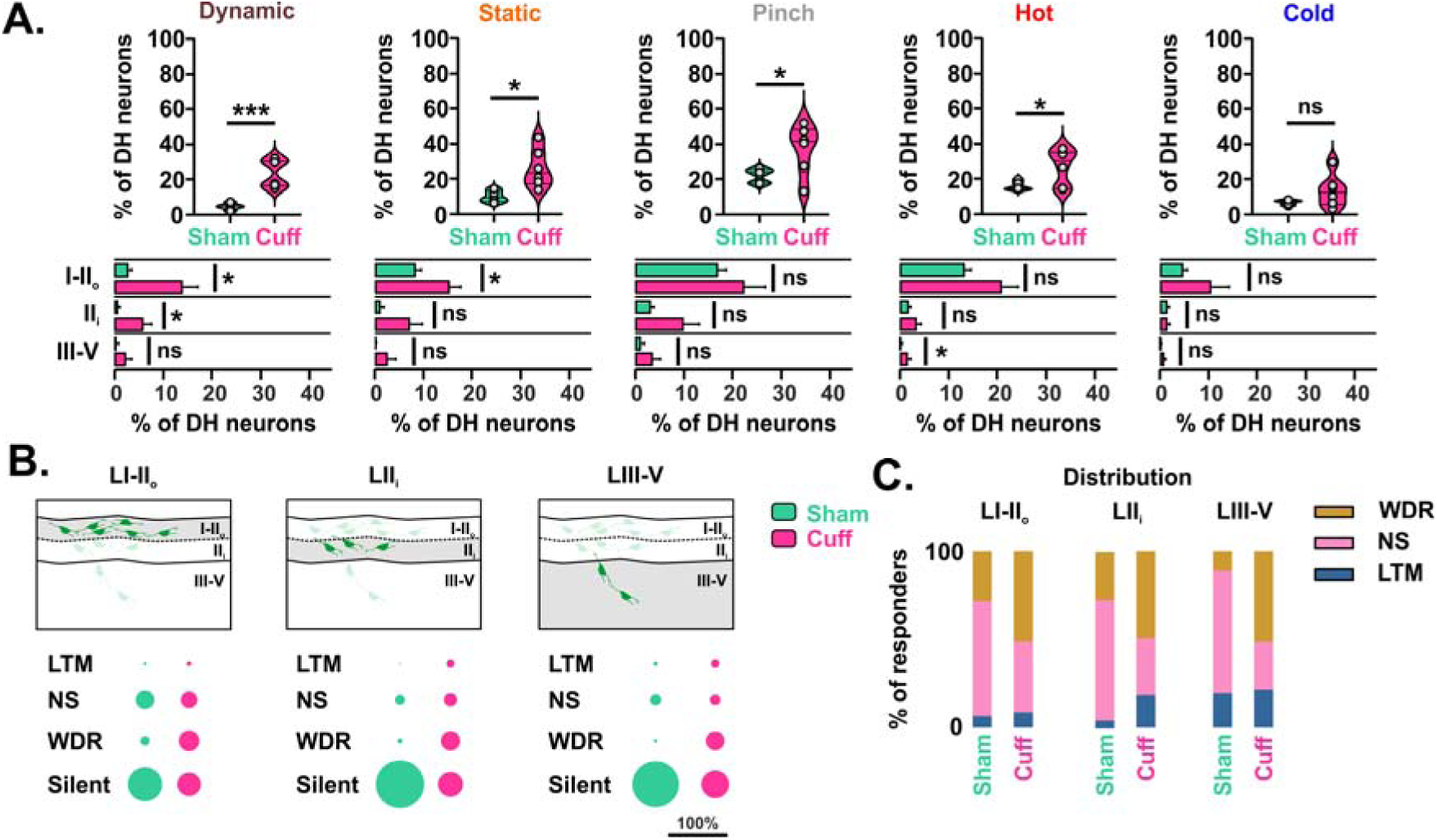
Nerve injury partially recapitulates the main features of spinal disinhibition, and selectively leads to hyperactivity of DH neurons following innocuous mechanical stimuli in lamina I-II_i_. **(A)** Percentage of DH neurons responding to physiological stimulations of the skin in Sham vs Cuff animals. Distributions of these neurons across laminae I-II_o_, II_i_ or III-V are indicated below each graph. Sham n=5, Cuff n=6 mice. ns= not significant, *p < 0.05, ***p < 0.001, unpaired t tests and Mann-Whitney tests. **(B)** Proportions of LTM, NS, WDR and silent neurons in Sham vs Cuff animals within laminae I-II_o_, II_i_ or III-V. **(C)** Distribution of responding neurons in Sham vs Cuff animals within laminae I-II_o_, II_i_ or III-V.

We then assessed if sensory processing through DH neurons was modified after nerve injury. Similar to what we observed following spinal disinhibition (Fig. S5A), the absolute number of NS neurons was relatively unchanged in neuropathic animals compared to shams, while the number of WDR and silent neurons drastically increased and decreased, respectively (Fig. S8B). These changes were again particularly pronounced in the deep DH (Fig. 6B). Consequently, NS neurons, which represented 66% of all responding neurons in sham animals, represented only 37% of all responding neurons after nerve injury (Fig. S8C). In contrast, about 50% of responding neurons were WDR neurons in each lamina after nerve injury, compared to 12-29% in shams (Fig. 6C). The number of LTM neurons was also affected by nerve injury but nearly exclusively in lamina II_i_, in which they made up about 20% of responding neurons, compared to less than 5% in shams (Fig. 6C).

Altogether, these data indicate that nerve injury sensitized DH neurons predominantly to innocuous mechanical stimuli, primarily through a decrease in the number of silent neurons and an increase in the number of WDR neurons in all laminae, and an increase in the number of LTM in lamina II_i_.

### Generation of a computational model of the plasticity of DH sensory processing associated with change in inhibition

In contrast to experiments using pharmacological disinhibition (Figs. 2-4), in which responses of each individual neuron could be compared before and after drugs application, we could not directly assess the plasticity of neurons recorded from neuropathic animals, from what they were before the injury (Figs. 5 and 6). However, given that data generated using spinal disinhibition revealed how sensory inputs are wired toward individual DH neurons, and how those are controlled by spinal inhibition (Fig. S7), we developed a computational model to determine the highest probability of plasticity in sensory processing that was induced by nerve injury, through spinal disinhibition. From 1000 generated DH models, we first selected the five best models that fit two main criteria: i) numbers of individual neurons responding to peripheral stimuli that were the closest to the values obtained experimentally in control conditions and after spinal disinhibition, and ii) models with the lowest number of disinhibited synapses from control to nerve injury, reflecting the most energy efficient plasticity induced by the neuropathy (Fig. 7A). To assess whether the models could reliably reflect experimental observations, we first compared the percentage of cells responding to each individual stimulus given by these five models to values obtained experimentally. In each case, the total number of responding neurons did not differ significantly between computational and experimental values in control conditions, after spinal disinhibition or nerve injury (Fig. S9A). The sensory modality of individual neurons was also not significantly different between computational and experimental data within lamina I-II_o_ (Table 2), lamina II_i_ (Table 3) or laminae III-V (Table 4). Finally, the proportion of LTM, NS, WDR and silent DH neurons (Figs. 7B and S9B), and the distribution of LTM, NS, WDR among responding neurons (Figs. 7C and S9C), were also consistent with what we observed experimentally in each conditions.

**Figure 7.**
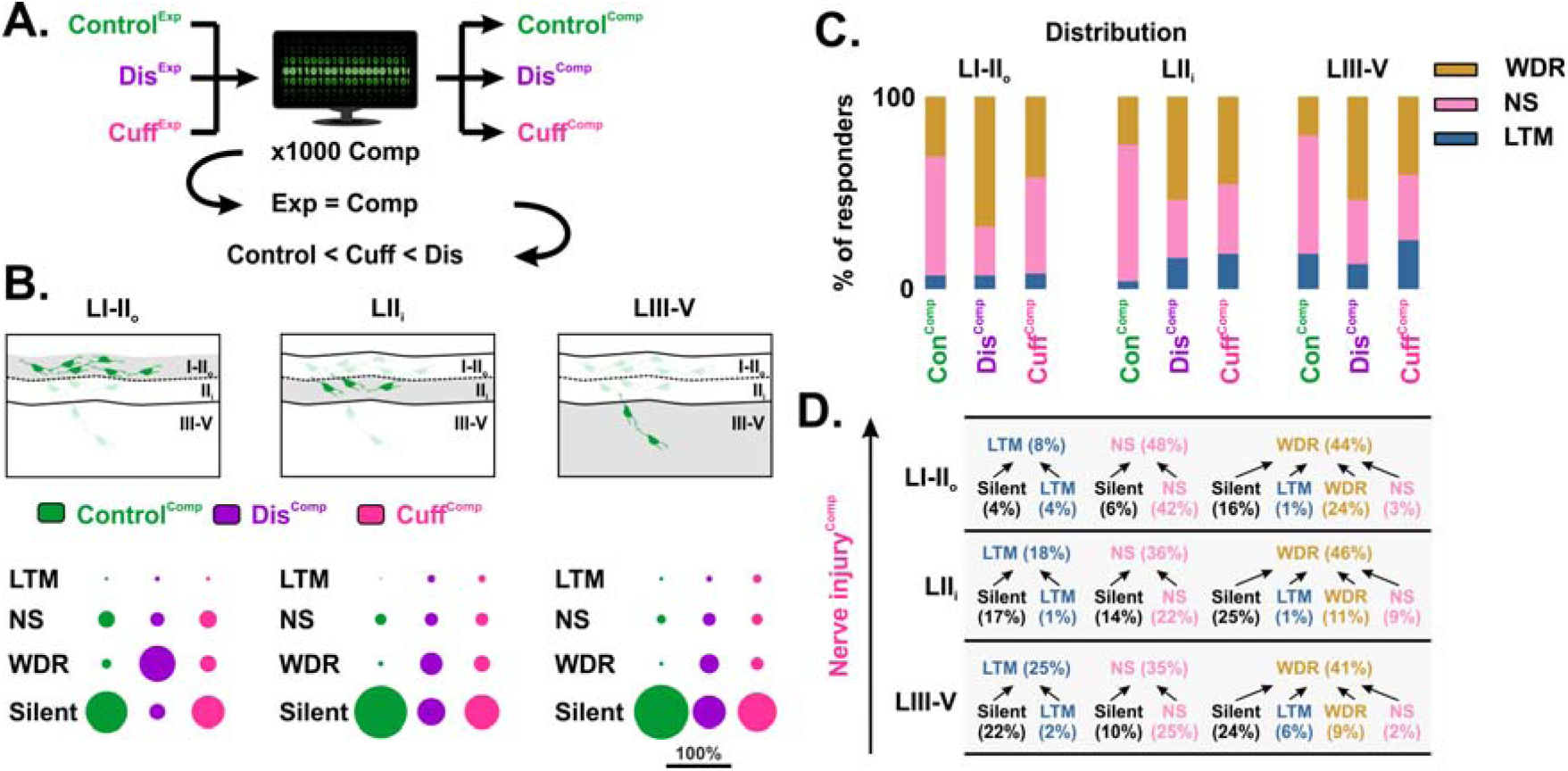
Computational modeling of the plasticity of DH neurons after nerve injury. **(A)** Schematic showing that experimental (Exp) data obtained using Ca^2+^ imaging in control mice (Con), after disinhibition (Dis), or nerve injury (Cuff), were used for computational modeling (Comp). Out of the 1000 models generated, the top 5% with computational values that were the closest to those obtained experimentally were first selected. Out of these models, the first 5 with the minimal number of mutations between control and neuropathic conditions were averaged to generate the final computational model of the DH. **(B)** Proportions of LTM, NS, WDR and silent neurons generated by the computational model in control conditions, after spinal disinhibition or nerve injury within laminae I-II_o_, II_i_ or III-V. **(C)** Distribution of responding neurons within laminae I-II_o_, II_i_ or III-V in control conditions (Con), after spinal disinhibition (Dis), or nerve injury (Cuff), obtained using the computational model. **(D)** Framework of the DH network showing the major transformations in sensory processing observed after nerve injury within laminae I-II_o_, II_i_ or III-V. Arrows point toward the proportions of LTM, NS, WDR neurons observed after nerve injury, and originate from what they were in control conditions, according to the computational model.

Altogether, this analysis indicates that the models can reliably reflect observations made experimentally. We thus averaged these five models to generate a representative computational model of the DH sensory processing, which was next used for further analysis.

### Nerve injury primarily leads to hypersensitivity of silent neurons rather than NS neurons

We used the computational model to estimate how DH sensory processing was affected by nerve injury, within each individual neurons. Similar to what we observed after spinal disinhibition, the major plasticity that was induced by nerve injury led to an increase in WDR neurons through the recruitment of previously silent neurons, within all laminae of the DH (Fig. 7D). Interestingly, the proportion of these Silent->WDR neurons was nearly identical after spinal disinhibition or nerve injury in lamina II_i_-V, but represented only half of what was observed after disinhibition in laminae I-II_o_ (Figs. 7D and S10), suggesting that loss of inhibition onto these cells is particularly pronounced in the deep DH. The second major plasticity revealed by the model was a substantial number of silent neurons that became LTM after injury. Those were found nearly exclusively from lamina II_i_ to lamina V (Figs. 7D and S10), strengthening the idea that nerve injury disinhibits a circuit of neurons, with a part that is located in the LTM recipient-zone (Abraira et al., 2017). Most importantly, the model also showed that the large majority of NS neurons remained NS after injury within each lamina of the DH (Figs. 7D and S10). This thus strongly suggests that hypersensitivity of silent neurons, rather than NS neurons, is the main driver of the sensitization of the DH to innocuous stimuli, and most likely of touch-induced pain, after nerve injury.

Altogether, these data show that nerve injury primarily leads to the recruitment of previously silent neurons, that then respond to innocuous mechanical stimuli in the deep DH, and to a wide range of sensory stimuli in all laminae.

## DISCUSSION

Sensory processing undergoes profound remodeling after nerve injury within the spinal DH, leading to abnormal persistent pain. The current study used 2-photon Ca^2+^ imaging in a newly adapted semi-intact *ex vivo* preparation allowing to selectively and simultaneously visualize the activity of excitatory neurons within all laminae of the DH, following physiological stimuli of the mouse hind paw. Combining experimental data and computational modelling in control and neuropathic mice, we showed that DH neurons receive a substantial number of sensory inputs that are strongly controlled by spinal inhibition. Not only does this inhibition defines the sensory modality of spinal DH neurons, it also prevents the activation of a countless number of neurons that are thus silent in physiological conditions. Under pathological conditions, when such inhibitory control is disturbed, DH neurons become hyperexcitable and process sensory inputs through highly polymodal neurons, mostly coming from the pool of previously silent neurons. Our revised description of the plasticity of DH neurons underlying neuropathic pain significantly advances current understanding of neuropathic pain processing. It introduces a novel framework for pathological sensory processing in the DH, offering a foundation to guide future research into the mechanisms and circuits underlying neuropathic pain.

### Technical considerations

The main objective of this study was to analyze selectively excitatory neurons at the circuit level across the whole DH with minimal sampling bias, following peripheral stimulations similar to what was used for behavior analysis *in vivo*. To achieve this technical challenge, we developed a modified *ex vivo* preparation that conserved the neuronal connectivity from the skin to the spinal cord DH. While the influence from supraspinal areas, or other key factors such as circulating hormones or immune cells were absent in this preparation, we validated our approach by confirming nearly identical proportions of LTM, NS and WDR in physiological conditions compared to those reported *in vivo* using single unit electrophysiological recordings (Zhang et al., 2013).

We analyzed the activity of DH neurons using the ultra-sensitive GCamp6f calcium sensor, which is highly reliable to detect suprathreshold activity with at least 1 action potential (Chen et al., 2013). Most importantly, a clear single transient far above spontaneous baseline calcium fluctuations was evoked in individual DH neurons following as less as a single electric pulse from the lowest (25µA) to the highest (500µA) intensity used in this study. Because 0.1 ms electric pulses applied to the dorsal roots are known to normally evoke one single action potential in responding neurons (Duan et al., 2014; Lu et al., 2013; Zhang et al., 2018), this indicates that Gcamp6f is sensitive enough to detect DH neuronal activity following one single action potential, and that there is no floor effect in the sensitivity of capturing activated neurons using this approach (Fig S11A). However, sub-threshold variations of membrane potentials unable to elicit sufficient calcium increase to produce an action potential were most likely undetected under our imaging conditions. Therefore, neurons identified as “silent” in the current study may actually receive active afferent inputs from peripheral sensory neurons before disinhibition or nerve injury, which would have been measured as small excitatory postsynaptic potentials (EPSP) using more sensitive electrophysiological approaches. However, as these subthreshold inputs would not be sufficient to produce an action potential, which would otherwise have been detected by Gcamp6f, no release of neurotransmitters would be evoked. Such neurons would then still be considered functionally “silent”, or “gated” as described in other studies, at the circuit level.

While the kinetic of Gcamp6f can detect multiple action potentials when fast laser scanning is performed continuously at the single cell level (Fig. S11), we could not report firing frequencies with our experimental protocol due to the low speed of laser scanning of the whole DH. To compensate for the low speed of laser scanning compared to fast calcium transients, we recorded during 10 seconds with repeated stimulations at 2Hz, a frequency that is known to reliably evoke action potentials in DH post-synaptic neurons without synaptic failure, even for neurons receiving C-fiber inputs (Dickie et al., 2017). At higher frequencies, calcium transient pics still reliably report each evoked electric pulses up to 2Hz (Figs. S11B and C). Peripheral stimulations at 2Hz resulted in a sustained 10 seconds long transient with an amplitude well above baseline, indicating that an increase in fluorescence from a responding neuron would be visible during the entire stimulation period in our protocol. Together, this indicates that virtually all activated cells of the imaging field could be systematically detected during the acquisition, even with slow laser scanning.

### A new perspective of the role of deep DH neurons in pain processing

Current views of the DH network rely on the existence of sensory modules organized within the layers of the DH that process different aspects of the somatosensory system (Gatto et al., 2021; Peirs et al., 2020). Consistent with previous studies, we found that in physiological conditions, the majority of neurons in the most superficial laminae are NS, which have been repetitively reported to be enriched in this area (Greenspon et al., 2019; Harding et al., 2020; Iseppon et al., 2022; Light and Lee, 2008; Nishida et al., 2014; Sullivan and Sdrulla, 2022; Zhang et al., 2013). Strikingly, using an unbiased sampling of excitatory neurons for the first time in the deep DH, brought by the newly adapted *ex vivo* preparation, we found that such enrichment in NS is also profound in excitatory neurons of deeper laminae in the absence of injury. This previously unappreciated proportion of NS neurons in the deep DH differs from previous studies reporting a greater number of WDR neurons in laminae III-V (Medrano et al., 2016; Zhang et al., 2013). However, it is important to note that previous studies describing the sensory modalities of deep DH neurons have mostly been performed using blind electrophysiological recordings, which typically used repeated mechanical probing of the skin to identify neurons of the receptive field. Given that i) the excitatory or inhibitory nature of WDR neurons remains to date relatively uncertain (Zhang et al., 2024b), ii) that it was recently shown that inhibitory neurons are much more sensitive to innocuous mechanical stimuli than excitatory neurons (Sullivan and Sdrulla, 2022), and iii) that the deep DH contains a high number of Pax2 positive neurons with large somas (personal communication), it is then possible that NS neurons are in fact more prevalent among excitatory neurons, which were selectively targeted in our study. It seems thus critical to characterize precisely the phenotype of WDR neurons of the deep DH in further functional investigations, as changes in the excitability of WDR neurons reported in previous studied will likely have functional opposite effects if they occurred in inhibitory, rather than excitatory, neurons. Nevertheless, the high number of NS neurons that we found in the deep DH coincides well with studies from various species showing that the deep DH contains a large number of antenna-type neurons expressing the substance P, with dendrites extending to the superficial laminae and associated with bundles of CGRP positive axons, with minimal A-fiber inputs (Carlton et al., 1990; Hunt et al., 1981; Jakab et al., 1990; Kokai et al., 2022).

### Contribution of LTM, NS and WDR neurons to neuropathic pain

Using functional Ca^2+^ imaging for the first time in the deep DH, we detected LTM neurons preferentially located in laminae III-V, consistent with previous descriptions of LTM neurons in this area (Abraira et al., 2017; Chirila et al., 2022; Light and Lee, 2008). LTM neurons were also observed in neuropathic animals, but those originated almost entirely from previously silent neurons. Some of these cells likely include neurons expressing the gamma isoform of protein kinase C (PKCγ), which are enriched in lamina II_i_ and lamina III (Peirs et al., 2020), and are activated by innocuous mechanical stimuli only under pathological conditions or intense walking (Malmberg et al., 1997; Miraucourt et al., 2007; Neumann et al., 2008; Peirs et al., 2016). Accordingly, we recently reported that chemogenetic silencing of these cells did not change any mouse behaviors following a battery of innocuous and noxious stimuli under physiological conditions, but significantly reduced mechanical hypersensitivity induced by nerve injury (Peirs et al., 2021). These Silent->LTM neurons are found in the Excit-3 and Excit-4 populations described in transcriptional studies (Russ et al., 2021), and selectively receive sub-threshold A-fiber inputs (Abraira et al., 2017; Neumann et al., 2008; Peirs et al., 2014; Peirs et al., 2021). Those might also include neurons expressing the cholecystokinin (CCK), the transcription factor c-Maf, and/or neurons that transiently express the vesicular transporter 3 (tVGLUT3), which are all enriched in the deep DH. Indeed, these neurons also receive subthreshold A-fiber inputs that can evoke action potential firing only after nerve injury, and participate in neuropathic mechanical hypersensitivity, but not in physiological conditions (Cheng et al., 2017; Frezel et al., 2023; Peirs et al., 2021). Given i) the anatomical location of these Silent->LTM neurons in the LTM-recipient zone, ii) that their receive monosynaptic inputs from A-fibers and iii) that ablation or silencing of these neurons has been shown to strongly reduce mechanical hypersensitivity, they likely represent the first node that relays tactile inputs to other DH neurons after nerve injury.

Interestingly, while we showed that most neurons that are NS in physiological condition can become WDR after complete spinal disinhibition, we found that the vast majority of these cells continued to respond only to noxious stimuli after nerve injury. NS->WDR neurons, which have been observed in the current and previous studies using spinal disinhibition (Lavertu et al., 2014; Miraucourt et al., 2007), might then represent another form of plasticity of DH neurons that leads to pain hypersensitivity, but independently of mechanisms underlying persistent neuropathic pain. Indeed, we recently found that inflammation or nerve injury for instance can lead to very similar pain behaviors, including mechanical static and dynamic mechanical hypersensitivity, but that are mediated by distinct DH circuits, which are most likely sensitized through different mechanisms (Peirs et al., 2021). Accordingly, it was shown that neurons expressing NK1R in lamina I can be activated by polysynaptic activity driven by Aβ fibers after spinal disinhibition (Torsney and MacDermott, 2006), but by heterosynaptic facilitation of Aδ inputs after inflammation (Torsney, 2011), suggesting again that mechanisms and neuronal circuits involved in central sensitization closely depend on pain etiologies. As it has been reported that NS neurons expressing NK1R are sensitized to become WDR after intradermal injection of capsaicin (Miraucourt et al., 2009; Warwick et al., 2022), it is likely that NS->WDR plasticity relates more to the release of inflammatory mediators and the recruitment of immune cells during early inflammatory processing, which diminishes about a week after nerve injury (Gaudet et al., 2011), than to long term plasticity responsible for persistent neuropathic pain. Consistent with this hypothesis, it was recently reported that topical capsaicin sensitizes NS projection neurons of the anterolateral tract to become WDR, while neuropathy resulted in the persistent activation of a normally silent population of projection neurons (Yarmolinsky et al., 2025). Importantly, this would imply that NS->WDR plasticity, a core principle of the gate control theory of pain (Melzack and Wall, 1965), is not responsible for persistent mechanical neuropathic pain.

Using *in vivo* electrophysiological recording of DH projection neurons, Lavertu et al. found that neuropathic pain unmasks a new population of spinothalamic tract (STT) neurons, which likely drives pain hypersensitivity. As those shared electrophysiological characteristic similar to that of NS neurons but with lower threshold, the authors suggested that a subpopulation of STT-NS was probably altered by nerve injury, turning them into “WDR-like” neurons (Lavertu et al., 2014). Because our Ca^2+^ imaging data show that nerve injury primarily leads to the recruitment of previously silent neurons with functional characteristics of WDR neurons, it is likely that the new population of STT neurons falls in fact into the class of Silent->WDR neurons identified in the current and a recent study (Yarmolinsky et al., 2025), although our data could not distinguish interneurons from projection neurons. Silent->WDR neurons might belong to the Ex6CHRM3 population, which have recently been observed in the superficial DH after sensitization by capsaicin (Warwick et al., 2022). Of note, our analysis does not rule out that plasticity of physiological WDR neurons could also play an important role in neuropathic pain, through changes in action potential firing frequency for example, which is hardly detectable with slow laser scanning Ca^2+^ imaging. However, as there is currently little evidence that encoding of innocuous stimuli are affected in WDR neurons in neuropathic pain models (Zain and Bonin, 2019), it seems that plasticity of silent neurons, rather than changes in NS or WDR sensory processing, is what leads to DH hyperexcitability and eventually pain after nerve injury.

### Plasticity of silent neurons as the main driver of neuropathic pain

A major finding of our study is the disinhibition of silent neurons as a dominant mechanism of the sensory plasticity induced by nerve injury. While several other studies have shown that innocuous peripheral stimuli can unmask a DH polysynaptic circuit after spinal disinhibition or peripheral injury (Cheng et al., 2017; Duan et al., 2014; Huo et al., 2023; Miraucourt et al., 2007; Torsney and MacDermott, 2006), current hypothesis are mainly built on the idea that this polysynaptic circuit represent a maladaptive functional connection between LTM and normally NS neurons, capable of turning touch into pain. While we confirm that disinhibition can lead to NS -> WDR plasticity, our findings strongly suggest that central sensitization after nerve injury is mediated by different mechanisms, primarily driven by the unmasking of a new and previously silent neuronal circuit, including previously silent spinal projection neurons (Yarmolinsky et al., 2025). A striking observation about this population of silent neurons is that they did not fail to evoke any Ca^2+^ transient in physiological conditions due to the application of an inappropriate range of stimuli, as they also did not respond to suprathreshold electric stimulations of the dorsal roots, except after spinal disinhibition or nerve injury. Several mechanisms may have led to the disinhibition of these silent neurons. For instance, it has been reported that the number of synapses from inhibitory interneurons expressing parvalbumin (PV) onto PKCγ neurons are reduced after nerve injury, leading to *de novo* activation of these cells by innocuous stimuli, resulting in mechanical hypersensitivity (Petitjean et al., 2015). Reduction in chloride ion gradient (Coull et al., 2003), or in the tonic firing rate of DH inhibitory interneurons, including those expressing PV, dynorphin (Dyn) or the neuropeptide Y (Npy) (Ma et al., 2023; Rankin et al., 2024; Tashima et al., 2021; Zhang et al., 2018), may also reduce the strength of inhibition onto DH silent neurons, leading to an increase in their input sensitivity.

Interestingly, it was recently suggested that disinhibition of silent neurons also occurs after heat-burn induced inflammation, leading to thermal hyperalgesia (Barkai et al., 2023). This implies that plasticity of silent neurons might be a critical mechanism shared by other pathologies associated with pain hypersensitivity. Given the impressive proportion of silent neurons within all laminae of the DH, and how much they participate after injury, they might represent a large pool of available neurons that can be rapidly recruited to overcome inhibition and react against threats detected by the peripheral nervous system. As our data showed that nerve injury induced the disinhibition of only a subset of these silent neurons, particular attention should be focused in future studies to determine the phenotype of the specific silent neurons underlying neuropathic pain, and what unique mechanisms freed them but not others from their inhibitory controls.

## Conclusion

The current study combined computational modeling and selective Ca^2+^ imaging of DH excitatory neurons, using a newly adapted somatosensory preparation of the skin-DH, to reveal the plasticity of silent neurons as a major mechanism of pathological sensory processing during neuropathy. Although our computational model was restrained to focus on spinal disinhibition under neuropathic conditions, other mechanisms of central sensitization that have been well described could be implanted in the algorithm in future studies. Nevertheless, the data confirm a growing body of evidences suggesting that neuropathic pain cannot be solely explained by a crosstalk between LTM and NS circuits. Instead, this work points to a critical role of the plasticity of silent neurons within all DH laminae as being responsible for neuropathic mechanical hypersensitivity, and potentially other symptoms of chronic pain. Future investigations focusing on the pathophysiology of these cells will be important to understand the selective mechanisms that lead to the activation of this new neuronal circuit after nerve injury.

## CONTACT FOR REAGENT AND RESOURCE SHARING

Further information and requests for resources and reagents should be directed to and will be fulfilled by the Lead Contact, Cedric Peirs.

## RESOURCE AVAILABILITY

### Materials Availability

All unique reagents generated in this study are available from the Lead Contact with a Materials Transfer Agreement.

### Data and Code Availability

Data and code generated in this study are available from the Lead Contact upon reasonable request.

## EXPERIMENTAL PROCEDURES

### Animals

All animal procedures followed the ethical guidelines of the International Association for the Study of Pain and ethical guidelines of the Directive 2010/63/UE of the European Parliament and of the local ethical committee (authorization project 11837-2017101816028463V5), on the protection of animals used for scientific purpose.

Wild type (WT) Adult C57BL/6J mice used in the present study were purchased from Janvier labs (Saint Berthevin, France). VGLUT2^Cre^ mice (Vglut2-ires-cre) (RRID: IMSR_JAX:016963), R26^lsl-tdTomato^ (Ai14) (RRID: IMSR_JAX:007914) and R26^lsl-GCamp6f^ (Ai95) (RRID: IMSR_JAX:028865) mice were obtained from The Jackson Laboratory (Bar Harbor, ME, USA). PKCg^CreERT2^ mice were generously provided by Dr. David Ginty (Harvard University, Boston, USA) and Dr. Emmanuel Bourinet (Institut de Génomique Fonctionnelle, Montpellier). Double heterozygote VGLUT2^cre/+^; R26^lsl-GCamp6f/+^ mice were used for Ca^2+^ imaging. We also used double heterozygote VGLUT2^cre/+^; R26^lsl-tdTomato/+^ and PKCg^CreERT2/+^; R26^lsl-tdTomato/+^ mice. Both male and female mice (4- to 8-weeks old) were used for this study. No significant sex differences were observed in this study and data from each sex were pooled. Mice were kept on an inverted 12:12 light/dark cycle (light OFF 7am-7pm) in micro-isolator caging racks (Tecniplast) with food and water provided *ad libitum*.

### Induction of Cre^ERT2^

Tamoxifen (T5648; Sigma-Aldrich) was dissolved in absolute ethanol and corn oil (at a ratio of 1:9) and used at a final concentration of 10mg/ml. Induction of Cre was induced in PKCγ^CreERT2^ mice with intraperitoneal (*i.p*.) injections (200 μL) once per day for 4 days consecutively.

### Chronic constriction injury (Cuff)

Chronic constriction of the sciatic nerve was induced using the cuff model as previously described (Yalcin et al., 2014). Briefly, mice were anesthetized with 2.5% isoflurane and a 5 mm incision was made parallel to the femur of the left hind limb. The common branch of the sciatic nerve was exposed by separating the muscle layers using forceps. A sterile 2-mm long section of PE-20 tubing was placed around the nerve after gently stretching and hydrating it with saline (0.9% NaCl). Finally, the incision was closed with 4-0 nylon sutures. Following the surgery, mice were maintained on a heating pad until they recovered from anesthesia. For Sham mice, the sciatic nerve was similarly exposed but without the cuff emplacement. Behavior analysis was performed the day before and during 3 weeks post-surgery. Ca^2+^ imaging was performed 3 weeks post surgeries.

### Behavioral tests

For behavioral experiments, measurements were taken blinded to the experimental groups.

#### Paw Withdrawal Threshold to von Frey Filaments

static mechanical sensitivity was assessed as previously described (Peirs et al., 2015). Briefly, mice were habituated to transparent Plexiglas chambers on an elevated wire mesh table for at least 30 minutes for two days, and prior to testing. Assessments were performed using a set of calibrated von Frey monofilaments using the Up-Down method, starting with the 0.4 g filament (Chaplan et al., 1994). Each filament was gently applied to the plantar surface of the hind paw for 3 seconds or until a response such as a sharp withdrawal, shaking or licking of the limb was observed. Rearing or normal ambulation during filament application were not counted. Filaments were applied at five-minute intervals until the threshold was determined. The 50% paw withdraw threshold (PWT) was determined for each mouse on both hind paws.

#### Paw Withdrawal Score to a paint brush

dynamic mechanical sensitivity was assessed as previously described (Duan et al., 2014). Briefly, animals were habituated to transparent Plexiglas chambers on an elevated wire mesh table as described above. Tests were performed by lightly moving a 2 dalbe 200 Martre pure paint brush across the surface of the hind paw from heel to toe. The behavior of the mice was assessed through a scoring paradigm based on the mice response to brush stimulations, with a score of 0 for no reflex; 1 if the mouse moved or paid attention to the stimulus with no sign of pain; 2 if the mouse doubled tap or produced a brief lifting (∼1-2 seconds) of the stimulated paw, and 3 if the mouse showed extensive biting, licking, guarding or lifting (> 3seconds) of the stimulated paw. The application was repeated 5 times with a 5 minute interval between applications. The allodynic score was calculated for both hind paws of each mouse.

#### Pinprick test

noxious mechanical sensitivity was assessed as previously described (Peirs et al., 2015). Animals were acclimated to transparent Plexiglas chambers placed on a wire mesh table as described above. A small insect pin (tip diameter = 0.03 mm) was applied 10 times with a 5 minutes interval between applications on the plantar side of each hind paw. Care was taken to apply a minimal pressure without penetrating the skin. If the animal showed aversive behavior (lifting, shaking, licking of the paw) a positive response was recorded. A negative response was recorded if the animal showed none of these reactions within 1 second of application and a percentage of positive responses were determined. Paw withdraw frequency (PWF) was calculated for both hind paws by averaging the positive responses from each mouse.

#### Plantar heat test (Hargreave’s test)

animals were placed in an acrylic chamber on a heated (30°C) glass plate and acclimated to the test chamber for 30 minutes during 2 days and then for 30 minutes on the 3^rd^ day prior to testing. Using a plantar analgesia meter (IITC, 50% intensity), a radiant heat source of constant intensity was focused on the plantar surface of the hind paw and the latency of paw withdrawal measured. The heat source was stopped upon paw withdrawal with a cutoff of 20 seconds to avoid injury. Heat sensitivity test was repeated 3 times on each hind paw with a 5 minutes interval between tests and the results were averaged for each paw, as described previously (Peirs et al., 2021).

#### Cold immersion test

mice were habituated for two days, and measurements were done on the third day. Cold sensitivity was tested by immersing the ipsilateral hind paw in a temperature-controlled water bath at 5 °C until paw withdrawal was observed (cutoff time 30 seconds), as previously described (Lolignier et al., 2015). The latency of paw withdrawal was calculated by averaging 3 responses separated by at least a 3 minutes interval.

#### Dynamic hot and cold test

Mice were habituated to a thermal plate (Bioseb) set at 25°C for 5 minutes then a ramp of increasing (hot ramp; 25-50°C) or decreasing (cold ramp; 25-0°C) at a 1°C ⋅ minute rate, as previously described (Yalcin et al., 2009). The animal nocifensive behavior was monitored using a video camera and the paw withdrawal threshold temperature was calculated as the temperature that elicited a jump.

### Immunohistochemistry

Mice were deeply anesthetized by an intraperitoneal (i.p.) injection of a mixture of 100 mg/kg ketamine and 20 mg/kg xylazine, and perfused trans-cardially with heparinized saline solution (2.5 mL heparin/L) followed by 4% paraformaldehyde (PFA). The spinal cord and lumber DRGs were collected, then post-fixed in 4% PFA overnight, and cryoprotected in 30% sucrose diluted in phosphate-buffered saline (PBS). Spinal cords were cut transverse at 30µm with a cryostat (Microm HM 550) and kept free-floating in PBS with 0.05% sodium azide until used. DRGs were cut at 10µm with a cryostat (Microm HM 550) and sections were directly mounted on gelatin-coated slides. For fluorescent immunostaining of spinal cords, sections were first washed three times for 10 minutes with PBS, then blocked with 10% normal donkey serum (NDS) in PBS + 1% Triton-X100 (PBS-T) for 2 hours at room temperature (RT) before being incubated in primary antibodies diluted in PBS-T overnight at 4°C. For fluorescent immunostaining of DRGs, sections were first washed three times for 10 minutes with PBS, then blocked with 4% normal donkey serum (NDS) in PBS + 1% Triton-X100 (PBS-T) for 2 hours at RT before being incubated in primary antibodies diluted in PBS with 2% NDS and 0.2% Triton-X100 overnight at 4°C. In all cases, sections were then washed three times for 10 minutes in PBS and incubated in AlexaFluor secondary antibodies (Jackson ImmunoResearch) diluted 1:500 in PBS-T, for 2 hours at RT. Slices were finally washed three times for 10 minutes with PBS, mounted on gelatin-coated slides and cover-slipped with Fluoromount-G (00-4958-02, ThermoFisher).

The following primary antibodies were used for immunofluorescence staining at the following dilutions: anti-NeuN raised in Guinea Pig (1:1000, ABN90, Merck Millipore), anti-Pax2 raised in Goat (1:200, AF3364, R&D Systems), anti-NF200 raised in Mouse (1:200, N0142, Sigma), anti-CGRP raised in Rabbit (1:1000, T4032, BMA Biomedicals). We also used the conjugated IB4-488 (1:200, I21411, ThermoFisher Scientific).

The following secondary antibodies were used: DyLight 405 Donkey Anti-Mouse IgG (H+L) (1:500, 715-475-150, Jackson ImmunoResearch), Alexa Fluor® 488 Donkey Anti-Rabbit IgG (H+L) (1:400, 711-545-152, Jackson ImmunoResearch) Cy™3 Donkey Anti-Mouse IgG (H+L) (1:400, 715-165-150, Jackson ImmunoResearch), Cy™3 Donkey Anti-Goat IgG (H+L) (1:400, 705-165-147, Jackson ImmunoResearch), Cy™5 Donkey Anti-Guinea Pig IgG (H+L) (1:400, 706-175-148, Jackson ImmunoResearch) and Alexa 647 Donkey Anti-Rabbit IgG (H+L) (1:500, 711-605-152, Jackson ImmunoResearch).

### Confocal microscopy

Spinal cord sections were imaged with a motorized Zeiss Axio Observer.Z1 / 7 confocal microscope and the Zen blue software (Carl Zeiss). Sections were excited with 405nm, 561nm or 640nm laser sources and 400-450nm, 567-620nm or 660-700nm emission filter kits, respectively. In order to suppress emission crosstalk, the microscope was configured to acquire channels in a sequential mode. Z-stack images were acquired using a Plan-Apochromat 20x/0.8 dry objective and GaAsP-PMT detectors with a z-step of 0.68 µm at a resolution of 1024x1024 pixels.

### Wide field fluorescent microscopy

Images of dorsal root ganglions were obtained by epifluorescent microscopy with a motorized fluorescence microscope Axio Imager 2 (Zeiss) and the Zen blue software (Carl Zeiss). Lumbar DH slices were imaged using a colibri 7 light source, a Plan-Apochromat 20x/0,8 objective and the following filter sets from Zeiss: 96 HE BFP, 38 HE eGFP, 43 HE dsRed and 50 Cy5. Tile images were acquired using an Axiocam 705.

### Skin-nerve-spinal cord slice *ex vivo* somatosensory preparation

Mice were deeply anesthetized with an i.p. injection of 100 mg/kg ketamine and 20 mg/kg xylazine. The back of the mouse and the leg were shaved with an electric clipper and a dorsal laminectomy was performed. A block containing the spinal cord and the left leg was harvested and quickly transferred to a dissection dish perfused with an ice-cold (4°C) solution containing (in mM) 194 sucrose, 2.0 KCl, 7.0 MgCl_2_, 26 NaHCO_3_, 1.15 NaH_2_PO_4_, 11 D-glucose, and 0.5 CaCl_2_ (290-300 mOsm.kg^-1^), pH 7.4 bubbled with 95% O_2_ and 5% CO_2_. The skin of the plantar side of the hind paw was dissected from the paw and the tibial branch of the sciatic nerve was then dissected up to the spinal cord. The dura mater and pia were removed, and the spinal cord with the attached skin was secured onto an agar block using insect pins. The pins were placed carefully to stabilize the skin and the contralateral side of the spinal cord, with precautions taken to avoid damaging the regions of interest. Finally, a parasagittal slice (450-470 µm) of the DH was obtained using a vibroslice (VT1200 S; Leica Microsystèmes SAS, Nanterre, France). We also used 550 µm transverses slices of spinal cords with the L4 or L5 DRG still attached embedded into 3% low-melting agarose (Fisher Scientific 10377033) for electric stimulations of dorsal roots. In all cases, slices were left to recover for at least 1 hour in an artificial cerebrospinal fluid (aCSF) containing (in mM): 128 NaCl, 3 KCl, 2.5 CaCl_2_, 1.5 MgSO_4_, 0.6 NaH_2_PO_4_, 25 NaHCO_3_, 10 glucose (300-310 mOsm.kg^-1^), pH 7.4, bubbled with 95% O_2_ and 5% CO_2_. After recovery, the *ex vivo* preparation was transferred to a recording chamber (volume ≈3 or 33 ml for transverse or sagittal slices respectively) and continuously perfused at 3 or 5 ml/min with oxygenated aCSF at room temperature (≈23°C).

### Spinal cord disinhibition

GABAergic and glycinergic spinal cord disinhibition was performed as previously described using bath application of 10 μM bicuculline and 1 µM strychnine (Torsney and MacDermott, 2006). Recording began 30 minutes after the onset of drugs application, and during continuous perfusion with bicuculline and strychnine.

### Dorsal root electric stimulations

Dorsal roots were stimulated using a suction electrode and a A365 stimulus isolator (World Precision Instrument) as previously described (Peirs et al., 2015). Roots were stimulated at 2 Hz with a pulse duration of 0.1 ms. Afferent fibers were stimulated at different intensities to recruit Aβ, Aδ and C fiber with 25, 100 and 500 μA pulses, respectively. To identify the receptive fields of DRGs or the DH using the somatosensory *ex vivo* preparation (Figs. 1D and S1B), the tibial and sciatic nerves were electrically stimulated with 500 µA, 2 Hz and using a prolonged pulse duration of 5 ms, to ensure suprathreshold activation of all neurons.

### Physiological stimulations of the skin in the *ex vivo* somatosensory preparation

Experiments were performed in light shielded faraday cage in complete darkness. The skin was mounted, in a separate circular cavity, over a thin perforated metallic mesh to support the skin. The experimenter used a red light to visualize the skin and deliver the somatosensory stimulations. To avoid any external light arriving to the lens from the red light source, a custom-made black opaque plastic tube was placed on top of the recording chambre around the lens to shield the lens. This configuration would limit any external light other than the fluorescence emitted from the sample to reach the lens. Recordings where strong motion artifacts in the field of view during the stimulation periods were excluded. Peripheral cutaneous stimulations were delivered to the plantar surface of the left hind paw. For innocuous dynamic mechanical stimulation, a thin, soft paintbrush (Dalbe 200 Martre pure) was used to stroke the glabrous skin of the hind paw from heel to toe using the pattern described in Fig. S1D. The skin was stimulated 30 times at ≈0.5 Hz, following a tone. Each stroke covered one third of the plantar surface of the hind paw, such that three consecutive stimulations covered the entire paw. This stimulation pattern was then repeated ten times. Innocuous static stimulation was performed by applying a round blunt rubber surface (area = 12.6 mm²) with a force of 0.2 g / mm^2^, without inducing significant motion artifacts during acquisition. Stimuli were delivered to the skin at ∼0.5 Hz following a tone, using the pattern described in Fig. S1D to cover most of the plantar surface in a consistent manner.

For noxious mechanical stimulation, the skin was pinched using blunted forceps (Buerkle™ 5386-0102) following the pattern described in Fig. S1D, delivered at ∼0.5 Hz following a tone. Stimulations were applied as consistently as possible across the skin without causing damage or inducing strong motion artifacts (as described in Kim and Chisholm, 2021).

For thermal stimulation, hot or cold aCSF was perfused into a small microchamber containing the skin. The temperature of the solution in the chamber was continuously monitored using a Perfusion Temperature Controller (Scientifica), with the sensor placed near the skin without direct contact. For hot stimulation, the solution temperature was increased from 22°C to 50°C, while for cold stimulation it was decreased from 22°C to 5°C, each over a period of ∼30 s, as shown in Fig. S1D.

### Two-photon calcium imaging

Ca^2+^ imaging was performed using an upright Zeiss Axio Examiner Z1 microscope and the Zen black acquisition software (Carl Zeiss). Images were acquired using a Plan-Apochromat 20x/1.0 water-immersion objective for the DRG and DH, and a digital camera AxoCam MRm or a Multiphoton LSM 7 MP (Carl Zeiss) for wide field and multiphoton acquisitions, respectively. Slices were first visualized using combined infrared and differential interference prior to Ca^2+^ imaging with two-photon microscopy. For each slice, four individual focal planes of 425x425 µm window, separated by 15-20 µm intervals were illuminated with a sapphire Mai Tai laser (Spectra-Physics, USA) at 920 nm. The border of the scanning area was set near the dorsal border between the white mater and the grey matter of the DH to allow the visualization of DH neurons located in both superficial and deeper laminae. For electric stimulations, baseline fluorescence intensity was first acquired during 15 seconds, then the dorsal roots were stimulated for 10 seconds, and fluorescence was recorded for an additional 15 seconds. For physiological stimulations of the skin, baseline activity was recorded for 70 seconds, then cutaneous stimulations were applied for ≈60 seconds followed by another 70 seconds of recording.

### Calcium imaging analysis

At the end of each experiment, the maximum number of responding neurons was assessed following suprathreshold stimulations of the dorsal root with 500µA, under spinal disinhibition. Only neurons that responded to such stimulation were kept further for analysis and represented the total number of DH excitatory neurons. Images were exported into MATLAB (MathWorks) for analysis with a custom-made toolbox. Using this toolbox, images from each recording were x-y motion corrected, and recordings where a significant displacement in the Z leading to the loss of the focal point were excluded. In turn, the ΔF/F and standard deviation of fluorescence over time for each focal plane were analyzed individually, and ROI manually drawn around clearly identified cell bodies. ΔF/F was calculated as ΔF/F =(F(t) - F0)/F0 where F(t) is the fluorescent value at a given time and F0 as the mean fluorescence during baseline recording. Neurons were considered responders when the ΔF/F during the stimulation window exceeded 3 times the standard deviation of the mean fluorescence during baseline and with at least a peak amplitude 20% higher than any other peak during the baseline period.

Data was calculated either as percentage of total DH neurons responding to dorsal root stimulations at 500µA under spinal disinhibition, or as percentage of responders within laminae in control or pathological conditions, as indicated in figure legends. For segmentation of neurons receiving Aβ-fibers inputs, only cells that responded to all 25, 100 and 500 µA stimulations were kept for analysis. Similarly, only cells that responded to both 100 and 500 µA stimulations were considered for neurons receiving Aδ-fibers inputs. Finally, only cells that responded to 500 µA, but not 25 or 100 µA stimulations, were considered for neurons receiving C-fibers inputs.

### DH lamination

Very high precautions were used to separate DH laminae in this study. In a separate series of experiments, we first imaged spinal cord transverse and parasagittal slices of PKCγ^CreERT2/+^; R26^lsl-tdTomatofl/+^ mice using the exact same preparations and acquisition parameters described above for Ca^2+^ imaging. The border between the gray matter and the white mater was determined to set the beginning of lamina I (LI) using transmission light under 800 nm illumination. Then, slices were scanned at 1050 nm to visualize the PKCγ plexus and the upper and the lower borders of this plexus were carefully drawn to identify the borders of the inner lamina II (LII_i_). We precisely calculated the mean borders of II_i_ from 3 independent slices along the dorso-ventral, and rostro-caudal axis, and used these parameters to identify the estimated length of laminae I-II_o_, II_i_ and III-V in parasagittal and transverse slices, as described in Fig. S1C.

### Algorithmic inference of the DH plasticity in neuropathic condition

We used an algorithmic approach to determine the transformations from a control to a neuropathic DH network. The same process was applied independently but identically for all laminae of the DH (lamina I-II_o_, lamina II_i_ and lamina III-V):

We first classified cells recorded experimentally in control conditions and after disinhibition by their responsiveness to the 5 sensory modalities (Dynamic (D), Static (S), Pinch (P), Hot (H), Cold (C)). The modalities expressed by a cell before inhibition removal are called *effective* modalities, the modalities appearing after inhibition removal are called *potential* modalities and the modalities never expressed by the cell are called *absent* modalities. We assigned the value 1 to the effective modalities, 0 to the potential modalities and −1 to the absent modalities, following the order DSPHC for the modalities. For example, a cell expressing D, H in basal condition and D, S, P, H after inhibition removal is coded by [1, 0, 0, 1, - 1]. As another example, a cell that is Silent before disinhibition and expressed D,P,H after disinhibition is coded [0, −1, 0, 0, −1]. Then, we calculated the percentages of the different cell categories found in control conditions. For example, in lamina I-II_o_ 1.4% of the cells are coded by [0, 0, 0, 0, 0], 2.7 % by [0, 0, 0, 0, −1], 0.7% by [−1, 1, 0, 0, −1] etc… Next, we constructed a list of 1000 cells respecting these percentages, representing a simplified model of the DH network and taking into account an error margin of maximum 30%. The second step was to transform the control network into the one recorded in neuropathic animals. For this purpose, we used an algorithm only authorizing disinhibition (i.e, a potential modality turning into an effective modality). In the context of our coding, it means that we only authorized the transformations of 0’s into 1’s. For example, the algorithm could turn a [0, 0, 0, 0, 0] cell into a [1, 1, 1, 1, 1] leading to a decrease of the percentage of Silent cells and an increase of the percentage of DSPHC cells, as observed experimentally in neuropathic animals. Using the graph of all possible transformations of the control network, we modified it recursively, until the network displays the percentage of cells responsiveness found experimentally in neuropathic conditions. As several final networks satisfying observations made in neuropathic animals can be found, we performed multiple trials of our algorithm up to find 1000 correct plastic networks. We then selected the first 5% with values that were the closest to those recorded experimentally in control conditions and after nerve injury. Finally, out of this list of models, we selected and averaged the first 5 models with the minimal number of transformations, reflecting the theoretical minimal energy needed to induce spinal disinhibition in neuropathic conditions. The transformations between control and neuropathic networks presented in Figs. 7 and S10 represent the means of the transformations obtained from these 5 models, for each lamina.

### Statistical analysis

All statistical analyses were performed using GraphPad Prism 9.0. Values are presented as Median ± quartiles for violin plots, or Mean ± SEM otherwise. Statistical tests used for each figure is indicated in the respective legends. For experiments addressing the effect of DH disinhibition on neuronal activity, data were analyzed using paired t tests after passing the Kolmogorov-Smirnov normality test, or matched-paired Wilcoxon tests otherwise. Data comparing sham and neuropathic animals were analyzed using unpaired t tests after passing the Kolmogorov-Smirnov normality test, or Mann-Whitney tests otherwise. Behavior data were analyzed with two-way ANOVAs with repeated measures followed by Dunnet or Tukey multiple comparisons post hoc test for within and between groups analysis respectively. Data comparing computational and experimental values were analyzed with unpaired t tests, or fisher exact tests. Differences were considered significant if p < 0.05. Figures were made using CorelDRAW X7.

## SUPPLEMENTAL INFORMATION

Supplemental Information includes 4 tables and 11 figures.

## ACKNOWLEDGEMENTS

We thank A. Monteil and C. Lemmers from the Vectorology facility, PVM, Biocampus Montpellier, CNRS UMS3426, Anne-Marie Gaydier for administrative assistance and Cyril Rivat for critical comments on the manuscript. Images used for schematics were provided by Pixabay under the Creative Commons Zero (CC0) license. Funding was provided by the Institut National de la Santé et de la Recherche Médicale (INSERM), the University Clermont Auvergne, the French government IDEX-ISITE initiative 16-IDEX-0001 (CAP 20-25), the Fondation pour la Recherche Médicale and the Agence Nationale de la Recherche ANR-18-CE16-0002-01 and ANR-19-CE14-0033-01.

## AUTHOR CONTRIBUTIONS

AN, RD and CP conceived the study. AN, JM and CP performed calcium imaging. AN performed surgical procedures. AN, JM and CP performed behavioral analysis. AN, LPM, KH and CP performed IHC. AN, FG, MZ performed mathematical and computational analysis. AN, FG, MZ, LPM, CP performed formal analysis. RD and CP acquired fundings. AN and CP wrote the original manuscript with contributions from all the other authors.

## Data availability statement

The data that support the findings of this study are available from the corresponding author upon reasonable request.

