## Supplemental material for "Sensory plasticity of dorsal horn silent neurons: a critical mechanism for neuropathic pain"

**SUPPLEMENTAL DATA**

**SUPPLEMENTAL FIGURES, LEGENDS**

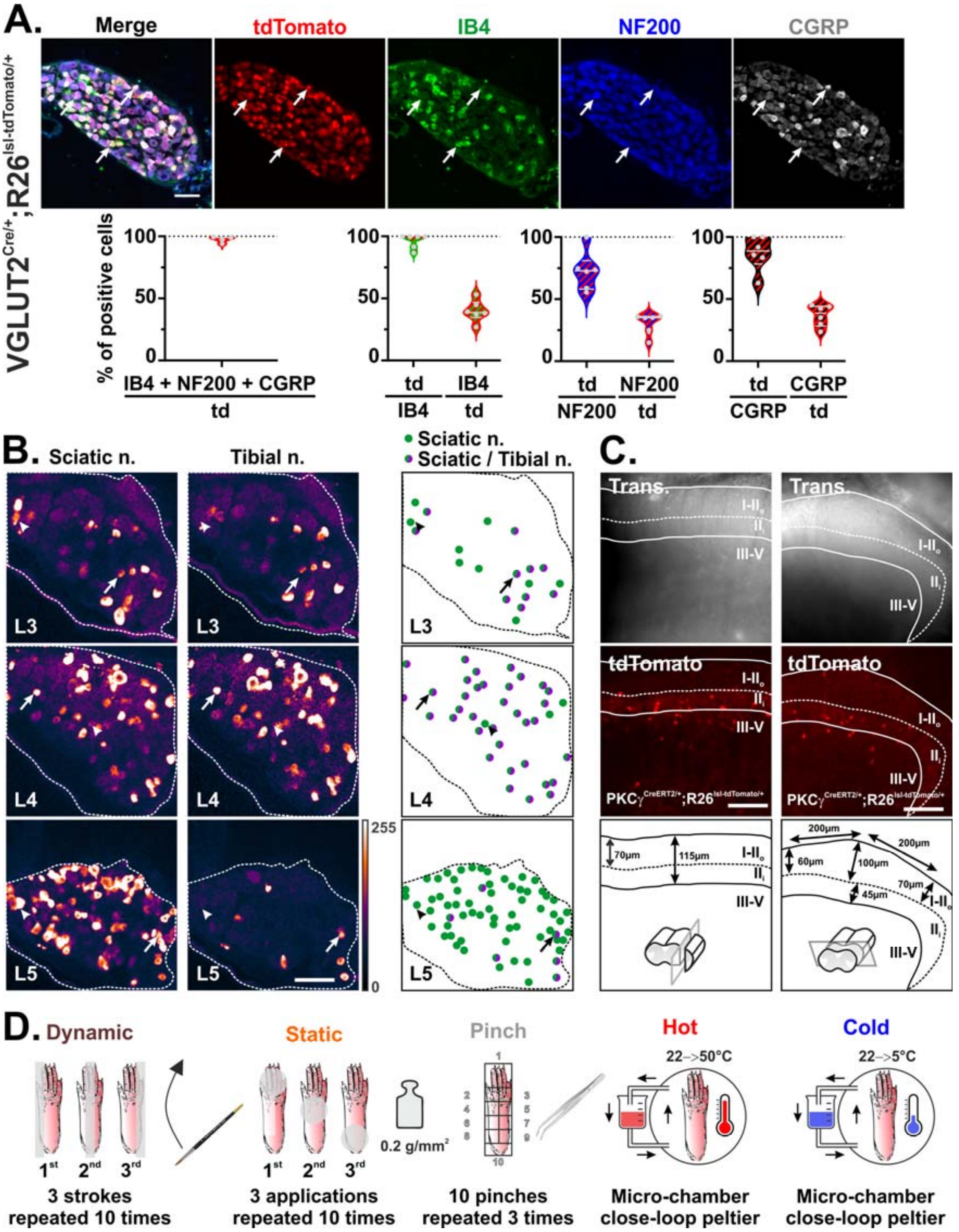


**Figure S1. Refers to Figure 1. Controls used to analyze neuronal activity at the population level.**

**(A)** Colocalization of virtually all tdTomato$^{+}$ neurons with IB4, NF200 and/or CGRP in VGLUT2$^{cre/+}$;R26$^{lsl\text{-}tdTomato/+}$ mice show that the transgene is expressed in myelinated, and in peptidergic and non-peptidergic unmyelinated, sensory neurons. About a third of tdTomato$^{+}$ neurons express IB4, NF200 and/or CGRP each, and most IB4, NF200 and/or CGRP express tdTomato, indicating that the majority of sensory neurons are represented through the expression of the VGLUT2$^{Cre}$ allele. Scale bar = 100 µm. **(B)** Representative images of Ca$^{2+}$ transients evoked in dorsal root ganglions at the level of L3, L4 or L5 in response to suprathreshold electric stimulation of the sciatic nerve (left) or its tibial branch (right). Right panels represent cells that are activated after sciatic (green dots), or both sciatic and tibial nerve (green / magenta dots) stimulations. Arrowheads and arrows show respective examples. Boundaries of laminae I-II$_{o}$ and III-V are indicated by continuous lines. Look up table (LUT) of pixel intensities is indicated at the bottom right. Scale bar = 100 µm. **(C)** 2-photon images of parasagittal (left) or transverse (right) slices of the spinal cord of PKCγ$^{CreERT2/+}$; R26$^{lsl\text{-}tdTomatofl/+}$ mice. Transmitted light was used to delineate the border of laminae I and the fluorescent plexus made by PKCγ neurons were used to delineate the borders of laminae II$_{i}$ as described in the method. **(D)** Schematic representing the stimulations paradigm used to stimulate the hindpaw with innocuous static and dynamic stimuli, and noxious mechanical hot and cold stimuli, using the *ex vivo* somatosensory preparation.

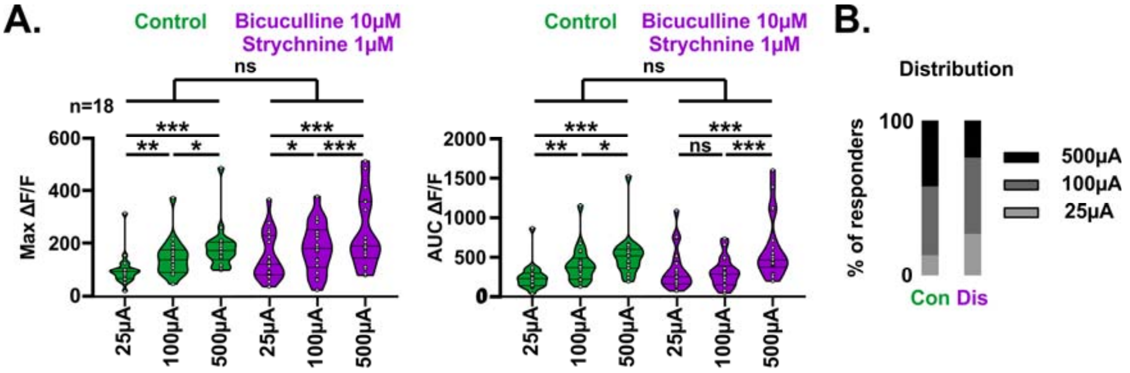


**Figure S2. Refers to Figure 2. Distribution of DH responsive neurons and calcium transients observed after peripheral electric stimulations before and after spinal disinhibition.**

**(A)** Maximum amplitude (left) and area under the curve (AUC, right) of $Ca^{2+}$ transients observed after stimulations at 25μA, 100μA and 500μA in control conditions (green) and after spinal disinhibition (magenta). n=18 cells. ns= not significant, *$p < 0.05$, **$p < 0.01$, ***$p < 0.001$, 2-way ANOVA with Tuckey's (for stimulation intensities) or Sidak's (for control vs disinhibition) post hoc tests **(B)** Distribution of responsive neurons following stimulations at 25μA (light grey), 100μA (dark grey) and 500μA (black) in control conditions and after spinal disinhibition. Con, control; Dis, disinhibition.

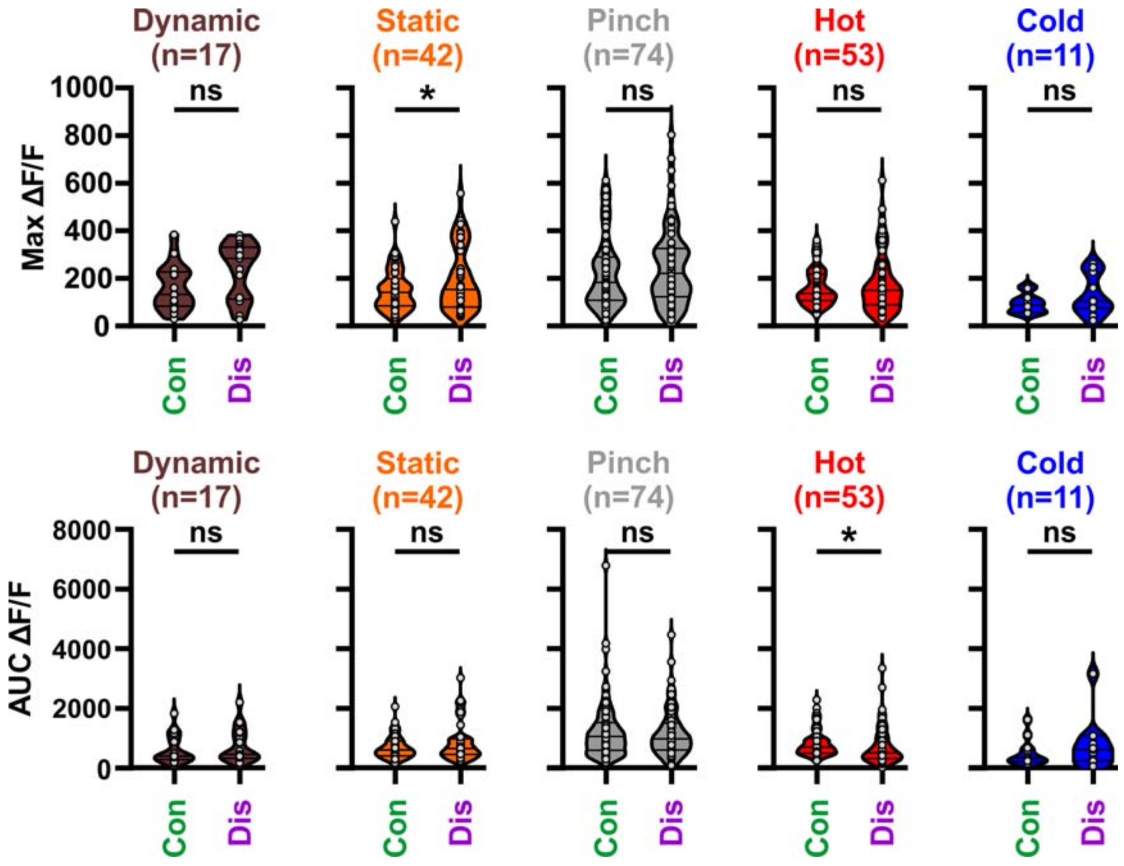


**Figure S3. Refers to Figure 3. Calcium transients observed after peripheral physiological stimulations before and after spinal disinhibition.**

The maximum amplitude (above) and area under the curve (AUC, below) of $Ca^{2+}$ transients following stimulations of the skin before (green) after (magenta) spinal disinhibition following innocuous dynamic and noxious mechanical stimuli, and noxious heat stimuli. The number of cells used for this analysis, before and after disinhibition, is indicated for each peripheral stimulus. Con, control; Dis, disinhibition. ns= not significant, *p < 0.05, paired t tests and Wilcoxon matched-pairs tests.

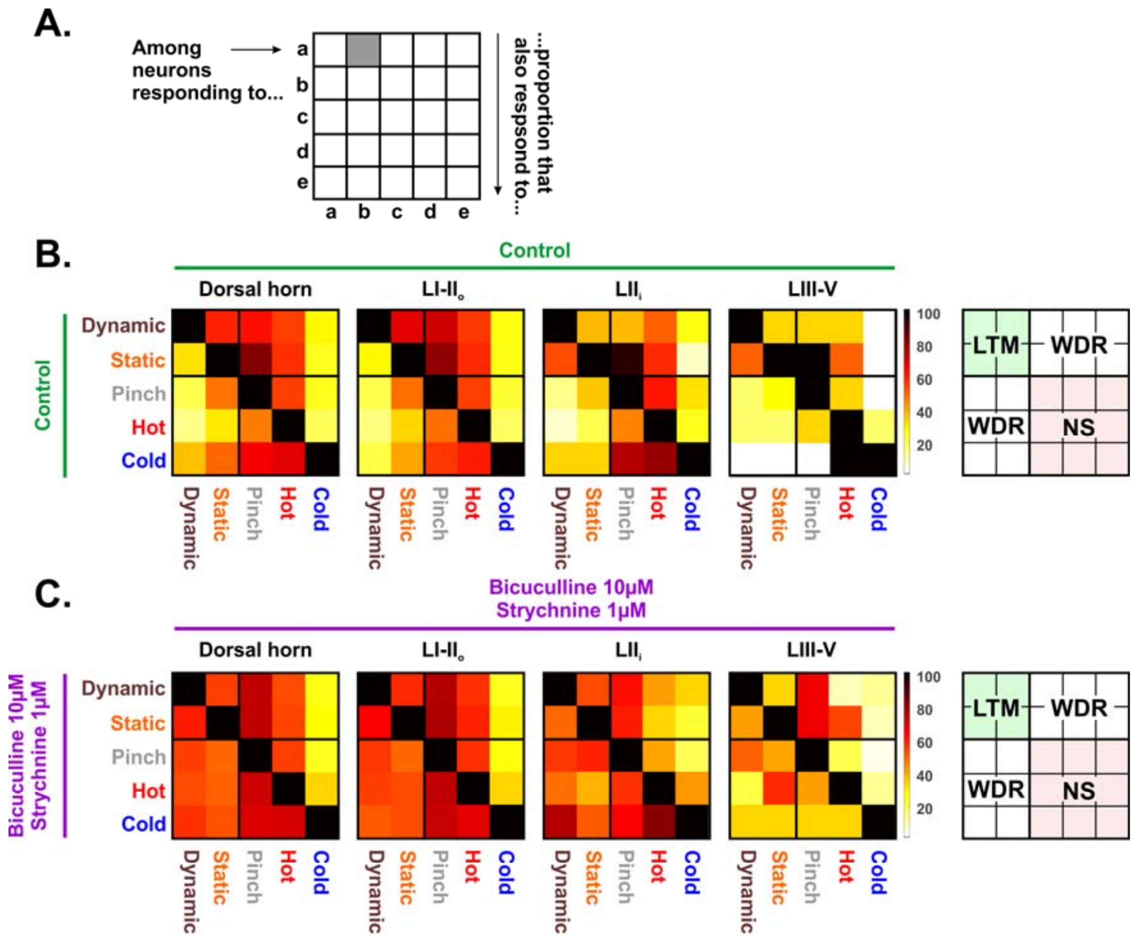


**Figure S4. Refers to Figure 4. Spinal disinhibition turns most DH neurons into polymodal cells.**

**(A)** Instructions on how to read heatmaps in B and C. Boxes within a line relate to cells responding to the modality indicated on the left of that line ("a", "b", "c", "d" or "e"). The value of each box represents the proportion of these cells that also respond to the modality of the related column, indicated below. For example, the box in grey represents the percentage of "a" cells that also respond to "b". **(B-C)** Heatmaps showing the sensory modality of neurons in control conditions **(B)**, and after spinal disinhibition **(C)**, on average and within laminae I-$II_o$, $II_i$ or III-V.

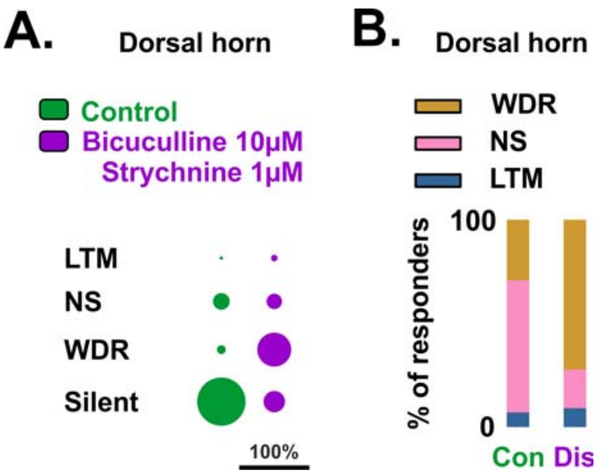


**Figure S5. Refers to Figure 4. Spinal disinhibition increases the number of wide dynamic range neurons and decreases the number of silent neurons.** **(A)** Proportions of LTM, NS, WDR and silent neurons on average in the DH in control conditions and after spinal disinhibition. **(B)** Distribution of responding neurons on average in the DH in control conditions and after spinal disinhibition. Con, control; Dis, disinhibition.

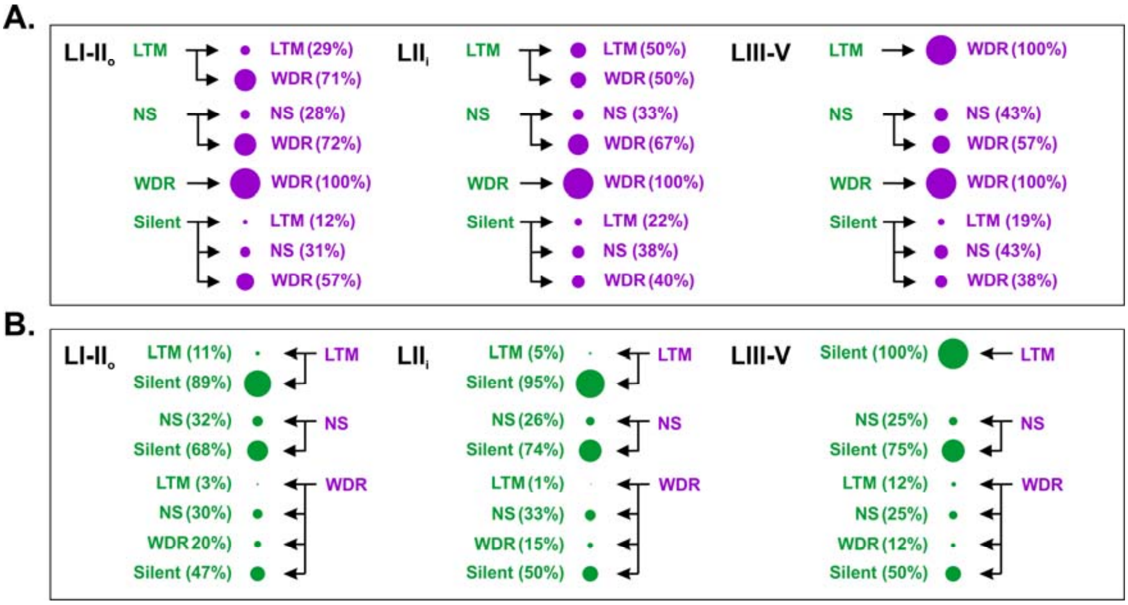


**Figure S6. Refers to Figure 4. Changes in sensory processing of LTM, NS, WDR and silent DH neurons after spinal disinhibition.**

**(A)** Proportions of LTM, NS, WDR and silent DH neurons observed in control conditions (green) normalized to what they became after spinal disinhibition, within laminae I-II$_o$, II$_i$ or III-V. **(B)** Proportions of LTM, NS, WDR DH neurons observed after spinal disinhibition (magenta) normalized to what they were in control conditions (green), within laminae I-II$_o$, II$_i$ or III-V.

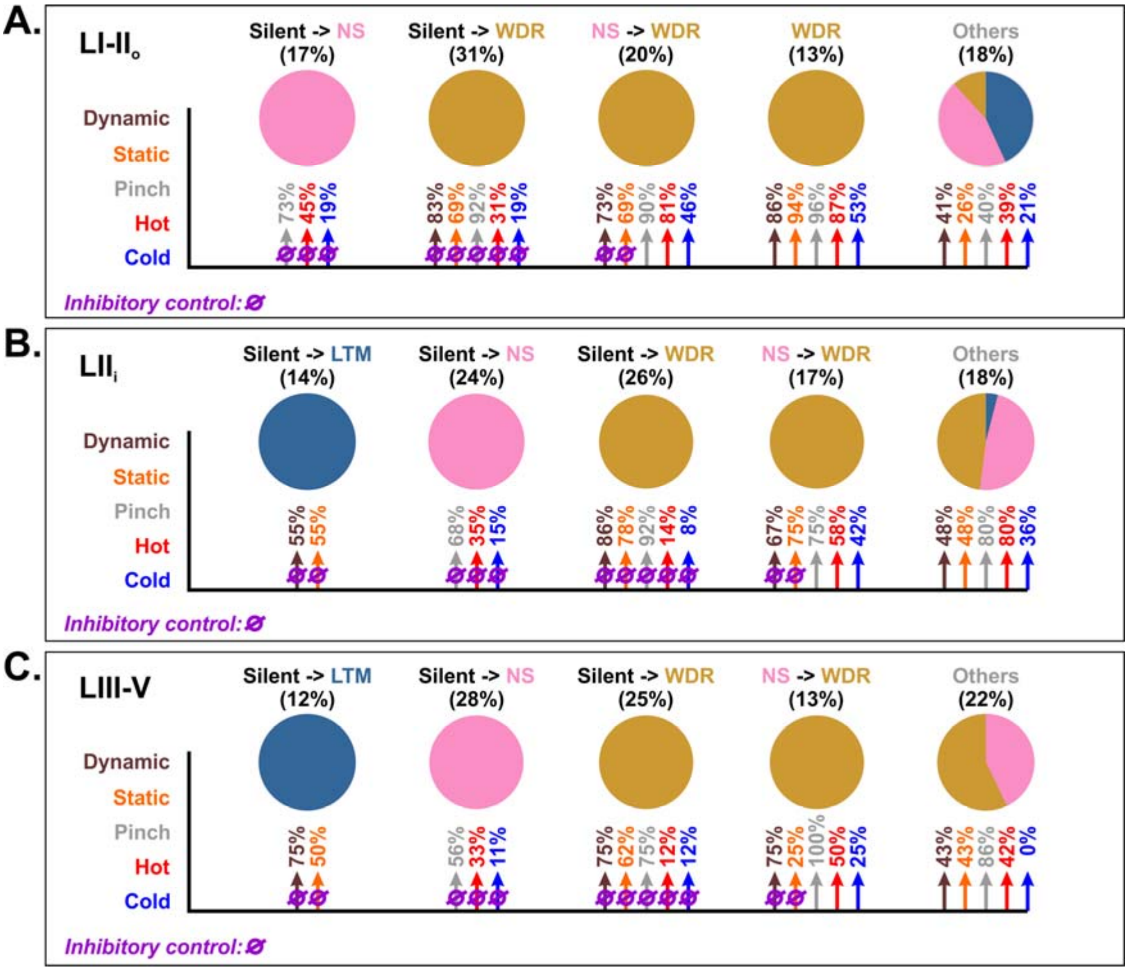


**Figure S7. Refers to Figure 4. Framework of sensory processing through spinal DH excitatory neurons.**

**(A)** Pie charts showing the main populations of neurons identified in laminae I-II$_o$, related to how they responded to peripheral stimulations in control conditions and after spinal disinhibition. Transitions from how these cells responded to peripheral stimulations in control conditions and after spinal disinhibition, and the proportion of these cells within this lamina, are indicated above the pie charts. For each pie chart, arrows and percentages below show their respective inputs from innocuous mechanical dynamic (brown) or static (orange) fibers, or noxious mechanical (grey), thermal heat (red) or cold (blue) fibers. Purple null symbols on top of theses arrows indicate that the corresponding input is silent under control conditions, but active after spinal disinhibition. **(B-C)** Similar framework of sensory processing through DH neurons within lamina $II_i$ and laminae III-V, respectively.

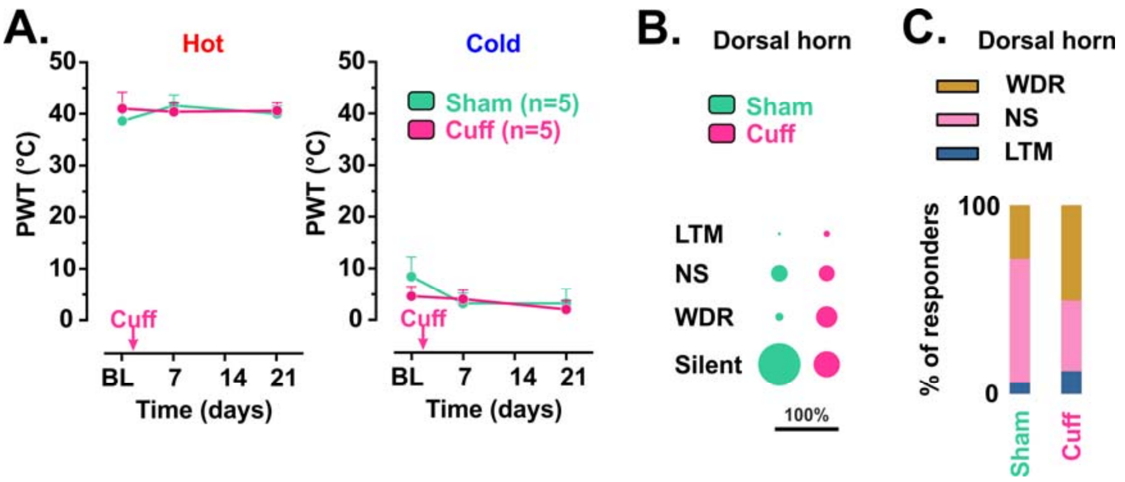


**Figure S8. Refers to Figures 5 and 6. Nerve injury does not affect thermal sensitivity, increases the number of wide dynamic range neurons and decreases the number of silent neurons.**

**(A)** Behavioral analysis of sensitivity to noxious thermal stimuli using the heat and cold plate tests, in Sham vs Cuff mice, in control conditions and up to 3 weeks after surgery. n=5 per group. 2-way ANOVA with repeated measures and Dunnett's (for comparisons within animal groups to baseline values) or Tuckey (for comparisons between Sham and Cuff values) post hoc tests. **(B)** Proportions of LTM, NS, WDR and silent neurons on average in the DH in Sham vs Cuff mice. **(C)** Distribution of responding neurons on average in the DH in Sham vs Cuff mice.

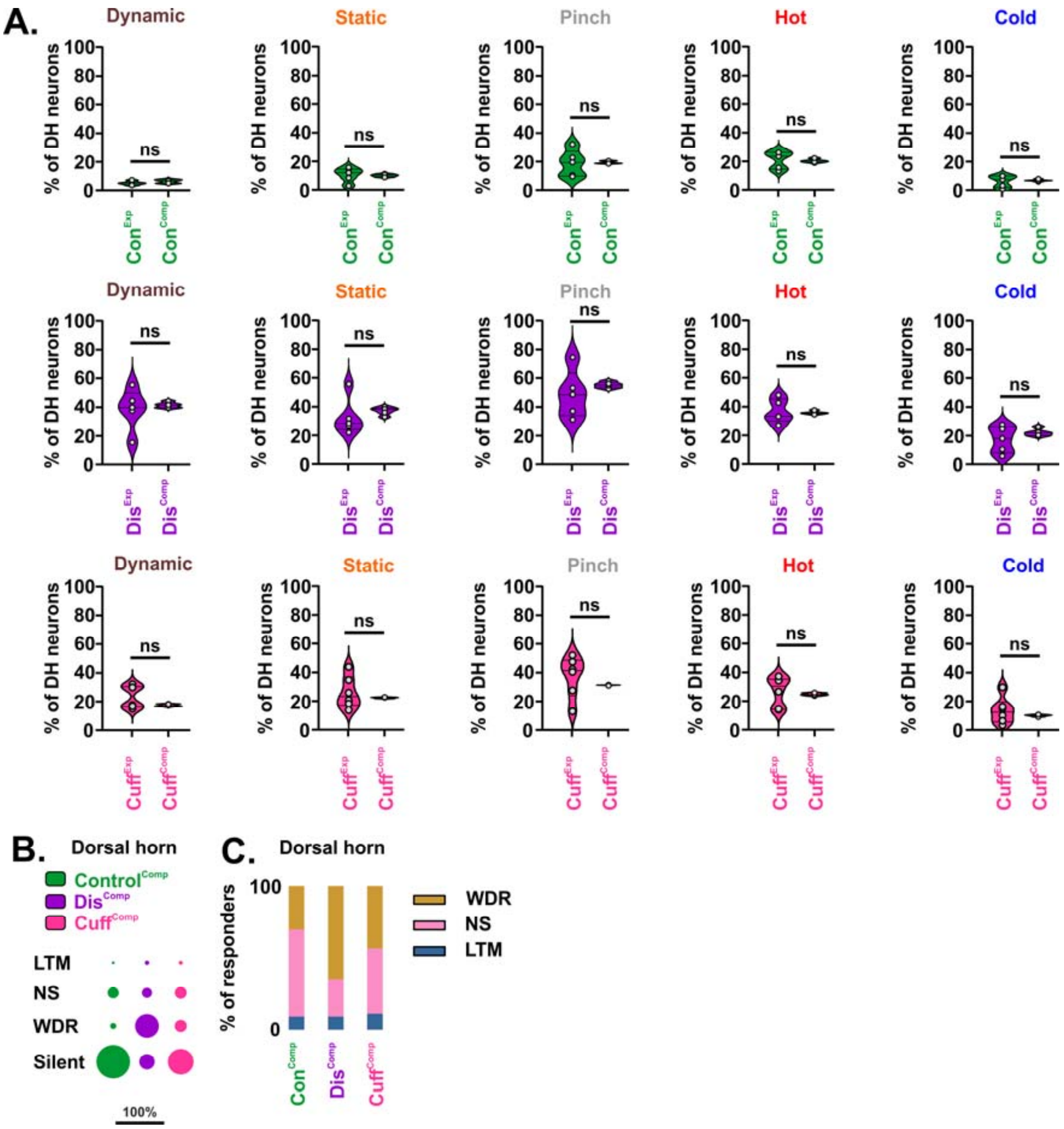


**Figure S9. Refers to Figure 7. Sensory modalities of DH neurons within laminae obtained using computational modeling.**

**(A)** Comparison of the percentages of DH neurons responding to physiological stimulations of the skin in control conditions (Con), after spinal disinhibition (Dis), or nerve injury (Cuff), obtained experimentally (Exp, as Figs 3C and 6A) or using computational modeling of the DH (Comp). n=5 mice or n=5 computational models. ns= not significant, unpaired t tests. **(B)** Proportions of LTM, NS, WDR and silent neurons on average in the DH in control conditions, after spinal disinhibition, or nerve injury using computational modeling. **(C)** Distribution of responding neurons in control mice, after spinal disinhibition, or nerve injury, within laminae I-II$_o$, II$_i$ or III-V using computational modeling.

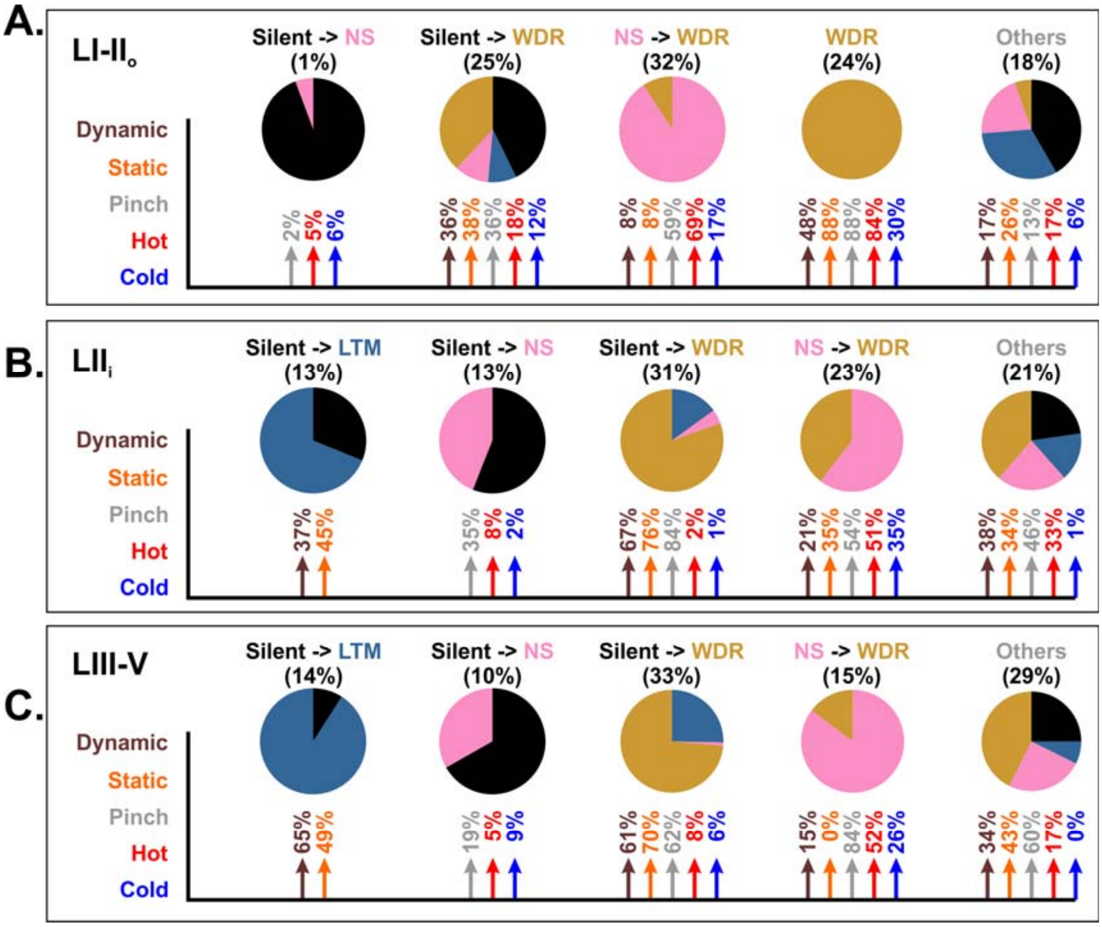


**Figure S10. Refers to Figure 7. Computational model shows the plasticity in sensory processing through spinal DH excitatory neurons after nerve injury.** **(A-C)** Pie charts showing the plasticity of the main populations of neurons identified in laminae I-II$_o$, lamina II$_i$ and laminae III-V, induced by nerve injury, according to the computational model. For each population, black, blue, pink and orange pies represent the proportion of cells that remained silent, or became LTM, NS or WDR after nerve injury, respectively. Proportion of these cells that are activated after nerve injury within each lamina, are indicated above the pie charts. For each pie chart, arrows and percentages below show their respective inputs from innocuous mechanical dynamic (brown) or static (orange) fibers, or noxious mechanical (grey), thermal heat (red) or cold (blue) fibers that are active after nerve injury.

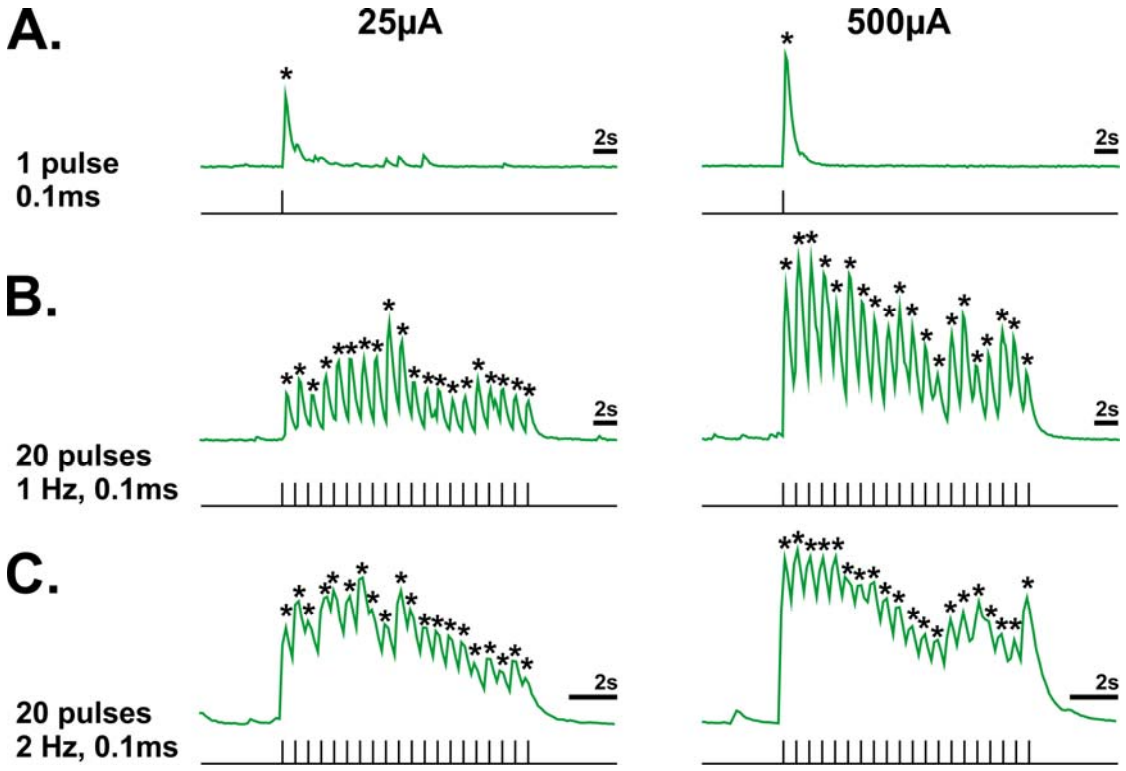


**Figure S11. Refers to Figures 1-6. GCamp6f calcium transients in DH neurons reliably detect peripheral sensory stimuli with low and high intensities, from one single electric pulse to 2 Hz trains.**

**(A)** Representative examples of calcium transients evoked by dorsal root electric stimulation at 25μA (left) and 500μA (right) with a single 0.1ms pulse, using fast laser scanning of a single dorsal horn neuron. **(B)** Calcium transients evoked in the same neuron following a train of 20 pulses at 1Hz. **(C)** Calcium transients evoked in the same neuron following a train of 20 pulses at 2Hz. For each panel, electric pulses are indicated in black below the green calcium transients. Transient pics indicated by stars show that GCamp6f reliably detects every electric pulses from one single pulse to 2Hz trains of stimuli at low and high intensities.

**Table 1. Analysis of NeuN$^+$ et Pax2$^+$ neurons in the dorsal horn of VGlut2$^{Cre/+}$; R26$^{lsl\text{-}tdTomato/+}$ mice.**

| | Td$^+$ (nb) | NeuN$^+$ (nb) | Pax2$^+$ (nb) | NeuN$^+$ that are Td$^+$ (%) | NeuN$^+$ that are Pax2$^+$ (%) | Td$^+$ that are Pax2$^+$ (%) | NeuN$^+$/Pax2$^-$ that are Td$^+$ (%) |
|---|---|---|---|---|---|---|---|
| Laminae I-IIo | 78±6 | 160±10 | 64±11 | 49%±5% | 40%±4% | 0%±0% | **89%±4%** |
| Lamina IIi | 55±11 | 137±15 | 54±11 | 61%±3% | 39%±3% | 0%±0% | **100%±1%** |
| Laminae III-IV | 34±3 | 87±5 | 34±3 | 56%±3% | 39%±1% | 0%±0% | **95%±2%** |

Mean ± SEM from 3 images for each animal (n=3).

**Table 2. Distribution of responding neurons in control conditions, after spinal disinhibition and in neuropathic animals, observed experimentally or after computational modeling, within laminae I-II$_o$**

| | $Con^{EXP}$ | $Con^{Comp}$ | *P* | $Dis^{Exp}$ | $Dis^{Comp}$ | *P* | $Cuff^{Exp}$ | $Cuff^{Comp}$ | *P* |
|---|---|---|---|---|---|---|---|---|---|
| LTM | 7.0% | 7.2% | >0.05 | 8.0% | 6.6% | >0.05 | 8.3% | 7.8% | >0.05 |
| $NS^M$ | 15.1% | 15.9% | >0.05 | 8.8% | 9.0% | >0.05 | 10.9% | 12.0% | >0.05 |
| $NS^T$ | 31.4% | 28.9% | >0.05 | 8.0% | 6.9% | >0.05 | 18.4% | 22.7% | >0.05 |
| $NS^{M/T}$ | 15.7% | 15.7% | >0.05 | 8.6% | 9.0% | >0.05 | 11.4% | 12.9% | >0.05 |
| $WDR^M$ | 6.4% | 5.8% | >0.05 | 23.5% | 20.8% | >0.05 | 14.2% | 12.8% | >0.05 |
| $WDR^{M/T}$ | 24.4% | 26.6% | >0.05 | 43.2% | 47.7% | >0.05 | 36.8% | 31.8% | >0.05 |

Distinctions between neurons that are mechano- (M), thermo- (T) or mechano- and thermo- (M/T) sensitive are indicated within each NS and WDR populations. *P* = Fisher's exact test between experimental (Exp) versus computational (Comp) matched values.

**Table 3. Distribution of responding neurons in control conditions, after spinal disinhibition and in neuropathic animals, observed experimentally or after computational modeling, within laminae $II_i$**

| | Con$^{EXP}$ | Con$^{Comp}$ | *P* | Dis$^{Exp}$ | Dis$^{Comp}$ | *P* | Cuff$^{Exp}$ | Cuff$^{Comp}$ | *P* |
|---|---|---|---|---|---|---|---|---|---|
| LTM | 4.8% | 3.7% | >0.05 | 15.0% | 15.6% | >0.05 | 17.9% | 18.0% | >0.05 |
| NS$^{M}$ | 19.4% | 20.5% | >0.05 | 12.9% | 12.5% | >0.05 | 17.3% | 15.7% | >0.05 |
| NS$^{T}$ | 30.6% | 36.1% | >0.05 | 10.2% | 10.0% | >0.05 | 12.3% | 15.9% | >0.05 |
| NS$^{M/T}$ | 17.7% | 14.1% | >0.05 | 8.8% | 7.6% | >0.05 | 3.1% | 4.3% | >0.05 |
| WDR$^{M}$ | 6.5% | 4.4% | >0.05 | 28.6% | 30.4% | >0.05 | 34.6% | 31.3% | >0.05 |
| WDR$^{M/T}$ | 21.0% | 21.3% | >0.05 | 24.5% | 23.9% | >0.05 | 14.8% | 14.8% | >0.05 |

Distinctions between neurons that are mechano- (M), thermo- (T) or mechano- and thermo- (M/T) sensitive are indicated within each NS and WDR populations. *P* = Fisher's exact test between experimental (Exp) versus computational (Comp) matched values.

**Table 4. Distribution of responding neurons in control conditions, after spinal disinhibition and in neuropathic animals, observed experimentally or after computational modeling, within laminae III-V**

| | Con$^{EXP}$ | Con$^{Comp}$ | *P* | Dis$^{Exp}$ | Dis$^{Comp}$ | *P* | Cuff$^{Exp}$ | Cuff$^{Comp}$ | *P* |
|---|---|---|---|---|---|---|---|---|---|
| LTM | 16.7% | 18.4% | >0.05 | 12.1% | 12.9% | >0.05 | 20.6% | 24.8% | >0.05 |
| NS$^{M}$ | 33.3% | 35.1% | >0.05 | 21.2% | 19.4% | >0.05 | 15.1% | 18.9% | >0.05 |
| NS$^{T}$ | 25.0% | 17.0% | >0.05 | 15.2% | 14.5% | >0.05 | 6.8% | 10.5% | >0.05 |
| NS$^{M/T}$ | 8.3% | 8.8% | >0.05 | 0.0% | 0.0% | >0.05 | 5.5% | 5.2% | >0.05 |
| WDR$^{M}$ | 8.3% | 13.1% | >0.05 | 33.3% | 33.2% | >0.05 | 35.6% | 33.4% | >0.05 |
| WDR$^{M/T}$ | 8.3% | 7.5% | >0.05 | 18.2% | 20.0% | >0.05 | 16.4% | 7.3% | >0.05 |

Distinctions between neurons that are mechano- (M), thermo- (T) or mechano- and thermo- (M/T) sensitive are indicated within each NS and WDR populations. *P* = Fisher's exact test between experimental (Exp) versus computational (Comp) matched values.
